# Electrical activity between skin cells regulates melanoma initiation

**DOI:** 10.1101/2021.12.19.473393

**Authors:** Mohita Tagore, Emiliano Hergenreder, Shruthy Suresh, Maayan Baron, Sarah C. Perlee, Stephanie Melendez, Travis J. Hollmann, Trey Ideker, Lorenz Studer, Richard M. White

## Abstract

Oncogenes can only initiate tumors in certain cellular contexts, which is referred to as oncogenic competence. In melanoma, whether cells in the microenvironment can endow such competence remains unclear. Using a combination of zebrafish transgenesis coupled with human tissues, we demonstrate that GABAergic signaling between keratinocytes and melanocytes promotes melanoma initiation by BRAF^V600E^. GABA is synthesized in melanoma cells, which then acts on GABA-A receptors on keratinocytes. Electron microscopy demonstrates synapse-like structures between keratinocytes and melanoma cells, and multi-electrode array analysis shows that GABA acts to inhibit electrical activity in melanoma/keratinocyte co-cultures. Genetic and pharmacologic perturbation of GABA synthesis abrogates melanoma initiation in vivo. These data suggest that electrical activity across the skin microenvironment determines the ability of oncogenes to initiate melanoma.

## Introduction

Melanoma arises at the dermal-epidermal junction, commonly harboring mutations in genes such as BRAF or NRAS^1^. These same mutations occur in benign nevi, raising the question of why some melanocytes, but not others, are competent to form melanoma. Previous work has shown that the developmental state of the cell plays a dominant role in such competence, since more neural crest-like melanocytes have a chromatin landscape that makes them permissive for oncogenesis^2^. Melanocytes in the skin are encased in a dense network of microenvironmental cells, including keratinocytes which make up the majority of the skin surface. We and others have previously shown that keratinocytes can have both pro-tumorigenic^3–5,8,9,10^ and anti-tumorigenic^6,7,11–13^ roles in melanoma. Whether these abundant microenvironmental cells like keratinocytes play a role in melanoma initiation remains unclear. Here, we identify a specific pro-tumorigenic keratinocyte population in direct communication with melanoma cells and identify the pathways mediating this communication.

## Results

### A reporter system to detect melanocyte/keratinocyte communication

In normal physiology, melanocytes are connected to keratinocytes through dendrites. These dendrites allow for the export of a pigment-containing organelle called a melanosome into the surrounding keratinocytes forming the epidermal melanin unit^14–16^. This unusual organelle transfer between cell types is responsible for skin coloration, since keratinocytes make up the vast majority of the skin surface. In addition to melanosomes, melanocytes also export smaller extracellular vesicles like exosomes that contain RNA and proteins^17^. We took advantage of this normal physiologic mechanism to develop a genetic reporter of melanocyte/keratinocyte communication. We engineered transgenic zebrafish to express Cre under the melanocyte-specific *mitfa* promoter, along with a floxed lacZ or GFP to RFP reporter under the keratinocyte-specific krt4 promoter. This approach has been previously used to study vesicular communication between different cell types, both during normal development as well as in cancer^18,19^. In this system, any keratinocyte which takes up *Cre* from the melanocyte will switch to express RFP fluorescence. To put this in the context of melanoma, we used the previously described miniCoopR transgenic system^20^, in which the melanocytes were further engineered to express *BRAF^V600E^* in the context of *p53*^−/−^ along with a palmitoylated GFP fluorophore (Fig. 1a). Upon transgene injection, control animals without Cre in the melanocytes had no RFP positive keratinocytes, as expected. In contrast, animals in which the melanocytes expressed Cre had on average 55% RFP positive switched keratinocytes (Fig. 1b and c). In 91% of the cases (Extended Data Fig. 1g), the RFP positive keratinocytes were directly adjacent to the *BRAF^V600E^, p53*^−/−^ melanocytes (marked by palmGFP fluorescence), suggesting it was only the subset of keratinocytes in physical contact with the nascent melanoma cells that exhibited such communication (Extended Data Fig. 1a - f).

**Fig. 1.**
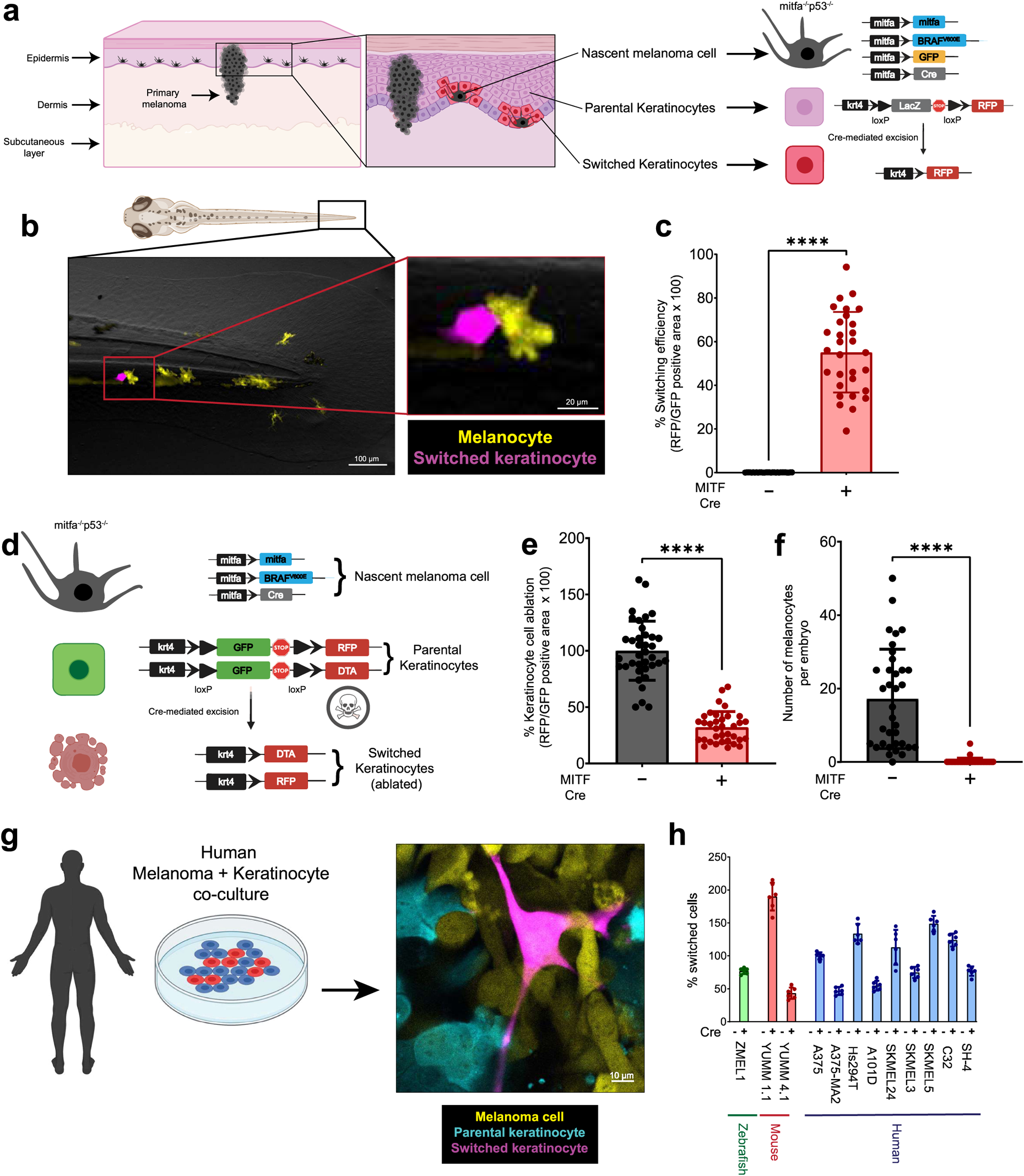
Nascent melanoma cells are in direct communication with keratinocytes. **(a):** Schematic representation of the genetic reporter system to identify keratinocytes in direct communication with nascent melanoma cells. (Left) The epidermal melanin unit is disrupted in primary melanoma, zoomed in image shows nascent melanoma cells and keratinocytes in direct physical contact. (Right) The genetic reporter system for detecting melanoma keratinocyte communication in zebrafish casper F0 embryos with the genotype *p53^−/−^ mitfa:BRAF^V600E^* injected with the indicated melanocyte and keratinocyte reporter constructs (+/− mitfa:Cre). **(b):** Representative image of an F0 zebrafish embryo with nascent melanoma cells over-expressing Cre and palmGFP in direct communication with a switched keratinocyte over-expressing RFP. Individual cells are pseudocolored as indicated. **(c):** % Switching efficiency calculated as % RFP positive area normalized to GFP positive area in 3 dpf zebrafish embryos. Data represent n = 30 control fish (negative for mitfa-Cre) and n = 30 switch fish (positive for mitfa-Cre) pooled from 3 biological replicates. Error bars: SD, P values generated by two tailed unpaired t test, **** p-value < 0.0001. **(d):** Schematic representation of the zebrafish genetic reporter system to detect melanoma keratinocyte communication and specifically ablate melanoma associated keratinocytes (RFP positive, switched) using the transgenic expression of DTA (diphtheria toxin gene A chain). **(e):** Switched keratinocyte ablation calculated as % RFP positive area normalized to GFP positive area in 3 dpf zebrafish embryos (+/− DTA expression) in keratinocytes. Absolute values were normalized to no DTA control to calculate cell ablation efficiency. Data represent n = 37 control fish (negative for mitfa-Cre) and n = 35 DTA fish (positive for mitfa-Cre) pooled from 3 biological replicates. Error bars: SD, P values generated by two tailed unpaired t test, **** p-value < 0.0001. **(f):** Number of pigmented melanocytes per embryo in +/− DTA conditions. Data represent n = 37 control fish (negative for mitfa-Cre) and n = 35 DTA fish (positive for mitfa-Cre) pooled from 3 biological replicates. Error bars: SD, P values generated by Mann Whitney test, **** p-value < 0.0001. **(g):** Representative confocal image of human melanoma/keratinocyte co-culture with non-switched keratinocytes, melanoma cells and switched keratinocytes pseudo-colored as indicated. **(h):** % Switching efficiency in human keratinocytes when co-cultured with zebrafish, mouse or human melanoma cell lines calculated as number of switched keratinocytes per well, normalized to the human melanoma cell line, A375. No switching was observed in the absence of Cre-expressing melanoma cell lines. Data represent n = 6 for control co-cultures (no Cre) and n = 6 for switched co-cultures (+ Cre) pooled from 3 biological replicates for each cell line indicated. Error bars: SD.

### Melanocyte/keratinocyte communication is required for melanoma initiation

This interaction between the keratinocytes and melanocytes could have been either pro or antitumorigenic. We tested this using a diphtheria toxin mediated cell ablation strategy previously developed in the zebrafish^21^. Using the same system as above, we re-engineered the keratinocyte cassette to express a floxed GFP to Diptheria toxin (DTA) transgene, such that any keratinocyte that receives Cre from the melanocyte would undergo cell death. This would help us understand if switched keratinocytes played any role in melanoma initiation, using our miniCoopR melanocyte rescue model. We also co-expressed a GFP to RFP switch cassette in the keratinocytes to mark the switched cells over time (Fig 1d). As a control, we found a significant decrease (68%) in the RFP positive area which labels switched keratinocytes in animals expressing the DTA cassette, validating that we could ablate keratinocytes in this setting (Fig. 1e). We then measured the number of pigmented BRAF^V600E^ positive melanocytes. While control animals not expressing keratinocyte-DTA had on average 17 rescued nascent melanoma cells, in the presence of keratinocyte-DTA this was reduced to an average of less than 1 (Fig. 1f). Because in the miniCoopR system, only these MITF positive pigmented melanocytes are capable of giving rise to melanomas that form later in life, this indicates that loss of communication with these keratinocytes is important for melanoma initiation by *BRAF^V600E^*.

### Human and mouse melanoma cells communicate with keratinocytes

To test the relevance of these findings in human and mouse melanoma progression, we developed a similar system using human or mouse melanoma cells in co-culture with human keratinocytes (Extended Data Fig. 2a). As a positive control we first engineered a zebrafish melanoma cell line (ZMEL1) to express Cre, and co-cultured it with HaCaT keratinocytes expressing a floxed dsRED to GFP reporter. Similar to what we saw in vivo in the zebrafish, we found that a consistent percent of cells (1.5% switching efficiency) in vitro also underwent Cre mediated switching. We then tested a panel of mouse (YUMM cells) and human lines (A375, Hs294T, etc) in the same fashion. While there was some variation from line to line, as expected, all of the tested lines exhibited Cre mediated fluorescent switching (Fig. 1g and h), and this again primarily occurred when melanoma cells and keratinocytes were physically adjacent to each other (Extended Data Fig 2d - i). To ensure that this was not a unique feature associated with HaCaT cells only, we tested this in a second human keratinocyte cell line, Ker-CT. Co-culture of Ker-CT keratinocytes with human melanoma cell lines showed similar robust Cre-mediated switching (Extended Data Fig. 3g), again only observed when melanoma cells and keratinocytes were physically touching each other (Extended Data Fig. 3c - f). Consistent with this, when the cells were separated by a Transwell membrane, the keratinocytes failed to undergo fluorescent switching, confirming that they require direct physical contact (Extended Data Fig. 2b and c). To eliminate the possibility of cell-cell fusion between melanoma cells and keratinocytes, resulting in the formation of GFP positive keratinocytes, we performed karyotypic analysis of individual cell populations as well as immunofluorescent studies for melanoma (SOX10) and keratinocyte (KRT14) markers. Karyotypic analysis of individual keratinocyte populations (dsRED positive and GFP positive) indicated that GFP positive keratinocytes were karyotypically identical to dsRED positive keratinocytes (Extended Data Fig. 3a). Further, we noted that GFP positive keratinocytes were negative for the melanoma marker, SOX10, thus eliminating the possibility of melanoma cell and keratinocyte fusion (Extended Data Fig. 3b). We also wanted to know if this form of communication was used by melanoma cells to communicate with each other (melanoma/melanoma crosstalk). To test this, we engineered a similar cassette as above with melanoma cells expressing a floxed dsRED to GFP switch reporter, and found no evidence of such communication between melanoma cells alone (Extended Data Fig. 3h - j). In addition, increasing the ratio of melanoma cells to keratinocytes increased switching efficiency suggesting that this mode of communication was dependent upon the density of melanoma cells (Extended Data Fig. 4a). Because the original studies using this Cre-based system was based on vesicle exchange, we wanted to confirm if this was the case here using both pharmacological and genetic loss of function approaches. Treatment of melanoma/keratinocyte co-cultures with GW4869 which is an inhibitor of exosome biogenesis, or genetic knockdown in melanoma cells of nSMase2 which is involved in exosome biogenesis^22^, substantially reduced keratinocyte switching efficiency (Extended Data Fig. 4b and c), highlighting the role of exosome-like vesicles in melanoma/keratinocyte communication. These data indicate that the vesicle mediated communication we report is a unique property only in the context of melanoma/keratinocyte communication across species.

**Fig. 2.**
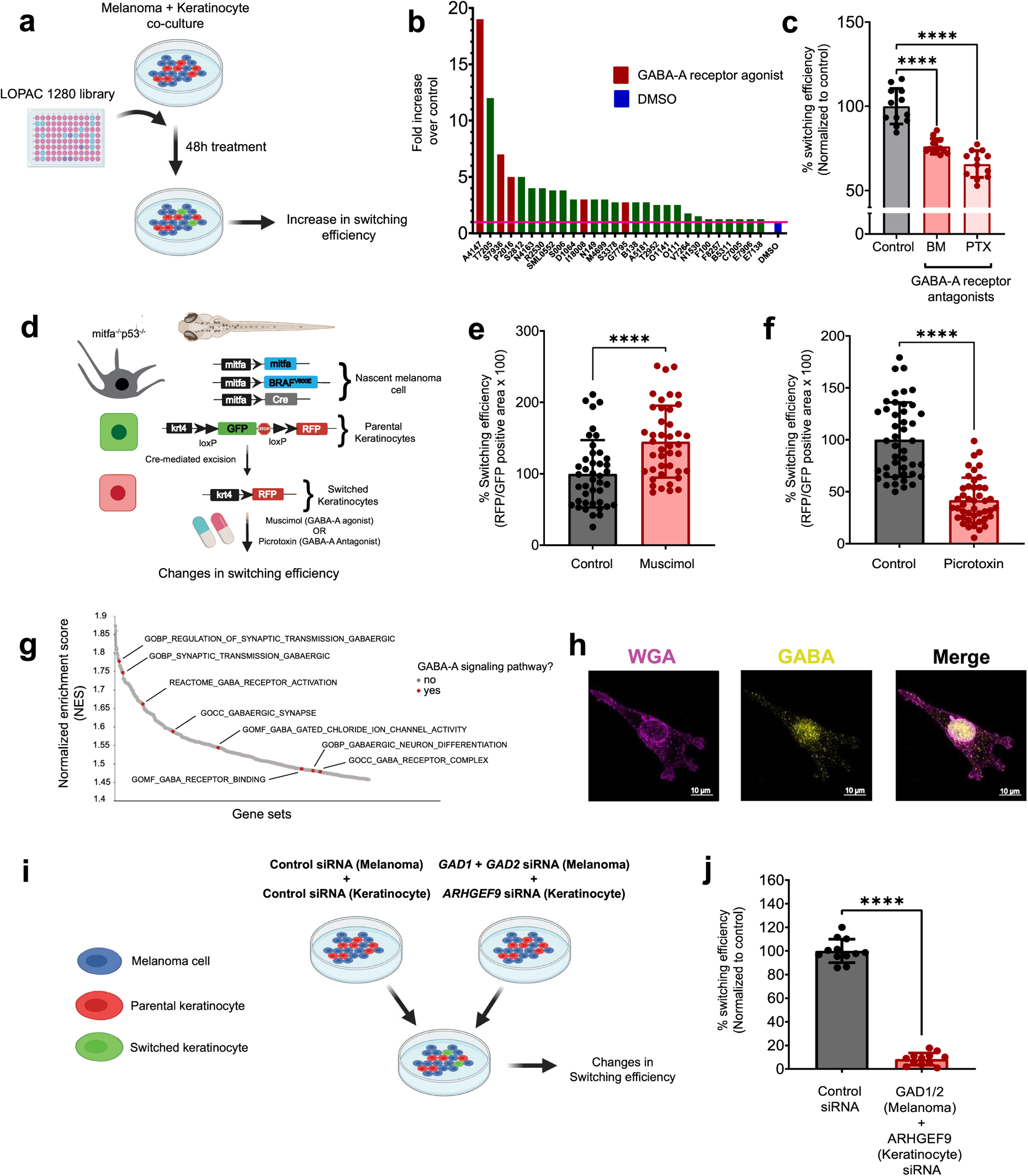
GABAergic signaling drives melanoma keratinocyte communication. **(a):** Schematic representation of the LOPAC small molecule library screen in human melanoma/keratinocyte co-cultures treated with control (DMSO) or 1280 LOPAC library compounds (10 µM each, indicated by their Sigma library identifiers) for 48 hours and quantified for increase in switching efficiency. **(b):** Fold change over control (DMSO) in switching efficiency in top 28 hits of the LOPAC small molecule library screen. Red bars indicate compounds which are agonists or allosteric modulators of the GABA-A receptor, blue bar represents DMSO control. **(c):** % Switching efficiency calculated as number of switched cells per well normalized to control (DMSO) upon treatment with GABA antagonists, BM (bicuculline methbromide, 100 µM) and PTX (Picrotoxin, 100 µM) in melanoma/keratinocyte co-cultures for 48 hours pooled from 4 biological replicates (n = 12). Error bars: SD, p-values generated by unpaired t test with **** = p < 0.0001. **(d):** Schematic representation of the F0 zebrafish genetic reporter assay to quantify changes in keratinocyte switching efficiency in zebrafish embryos treated with a GABA-A agonist (Muscimol) or a GABA-A antagonist (Picrotoxin). **(e-f):** % Switching efficiency calculated as % RFP positive area normalized to GFP positive area in 3 dpf zebrafish embryos treated with Muscimol (10 µM) or Picrotoxin (100 µM). Data represent n = 44 DMSO treated fish and n = 42 Muscimol treated fish and n = 44 Picrotoxin treated fish pooled from 3 biological replicates. Error bars: SD, P values generated by two tailed unpaired t test, **** p-value < 0.0001. **(g):** Waterfall plot of enriched pathways from GSEA analysis of switched vs parental keratinocytes. GABA receptor pathways are highlighted in red. **(h):** Immunostaining for GABA in A375 melanoma cells and membrane staining with wheat germ agglutinin (WGA). Individual cells are pseudocolored as indicated. **(i):** Schematic representation of the human in vitro switch reporter assay in melanoma keratinocyte co-cultures with genetic loss of function in GABA pathway components. **(j):** % Switching efficiency calculated as number of switched cells per well normalized to control siRNA when co-cultures are treated with a combination of GAD1/2 (melanoma cells) and ARHGEF9 (keratinocytes) targeting siRNA pooled from 3 biological replicates (n = 12). Error bars: SD, p-values generated by two-tailed unpaired t test with **** = p < 0.0001.

### A screen for melanoma/keratinocyte communication reveals a role for GABA

The mechanisms regulating this melanoma/keratinocyte crosstalk are unknown, but could represent a means for abrogating melanoma initiation. To address this, we performed a small molecule screen to identify pathways which mediated this communication. We used human A375 melanoma cells expressing Cre along with HaCaT keratinocytes expressing the floxed dsRED to GFP reporter, and then used fluorescent imaging to calculate the number of switched cells after applying the LOPAC1280 small molecule library, which contains a diverse set of chemicals affecting well-defined biological pathways (Fig 2a). Overall, we found 28 molecules which increased switching above the DMSO control wells (Supplementary Table 2). Amongst the top 10 hits, we found 3 molecules (30%) which were all involved in GABAergic signaling. For example, the top hit from the screen was the GABA-A receptor agonist, homotaurine (3-Amino-1-propanesulfonic acid sodium) which caused a nearly 20-fold increase in keratinocyte switching compared to control. Other hits included the GABA-A positive allosteric modulators SB205384 and tetrahydrodeoxycorticosterone (THDOC) (Fig 2b). To validate results from our screen, we tested switching efficiency in co-cultures with GABA, a natural agonist of GABA-A receptors and muscimol, a previously validated, ionotropic, GABA-A receptor agonist^23^ (Extended Data Fig. 5a), and found significant increases in switching efficiency in melanoma/keratinocyte co-cultures only (34% and 57% increase respectively) (Extended Data Fig. 5b). We further tested the GABA-A receptor antagonists bicuculline methbromide and picrotoxin, and found they significantly decreased Cre mediated recombination and switching efficiency in keratinocytes (24% and 35% decrease respectively), further confirming the role of GABA-A in this communication (Fig. 2c). To test whether GABAergic signaling mediated melanoma/keratinocyte communication in vivo, we used the transgenic zebrafish system described above to calculate switching efficiency (Fig 1c and 2d). We bathed the fish in either a GABA-A agonist (muscimol) or a GABA-A antagonist (picrotoxin) and measured the number of keratinocytes that had switched to RFP fluorescence (Fig 2d). Consistent with the in vitro results of the screen, we found that GABA-A agonist activity increased the number of RFP positive cells (44% increase, Fig 2e), whereas GABA-A antagonist activity strongly reduced the number of RFP positive cells (59% decrease, Fig 2f). These data support the notion that GABA is a specific mediator of melanoma/keratinocyte communication.

GABAergic genes are differentially expressed in keratinocytes versus melanoma cells GABAergic signaling has mainly been studied in the context of neuronal communication, but our data suggested it may unexpectedly play an analogous role in melanoma/keratinocyte communication. In neurons, GABA is synthesized in presynaptic neurons via GAD1 or GAD2^24,25^, and is exported into the synapse where it binds to GABA-A receptors on postsynaptic neurons. We hypothesized that components of this machinery might also be expressed by melanoma cells and keratinocytes, since previous studies have reported the presence of certain GABAergic signaling components in skin^26–29^. To test this, we performed RNA-sequencing of the keratinocyte populations that had undergone Cre mediated switching. Consistent with our chemical screen, Gene Set Enrichment Analysis demonstrated that those keratinocytes had a marked enrichment for pathways related to activation of the GABA-A receptor (Fig 2g, Supplementary Table 1), with upregulation of individual genes including GABA-A receptor subunits such as *GABRA3, GABRB3* and *GABRG2* as well as *ARHGEF9* (collybistin), a gene encoding an assembly protein which ensures proper synaptic organization of the GABA-A receptor^30–32^ (Extended Data Fig. 5c). We also analyzed publicly available gene expression data (from CCLE and Wistar Melanoma Cell lines)^33^ and found that melanoma cells (but not mature melanocytes) express high levels of the GABA synthesizing enzyme *GAD1* (Extended Data Fig. 5d). Moreover, *GAD1* expression is induced upon oncogene (*BRAF^V600E^*) expression and correlated with melanoma oncogenic competence in our previously developed hPSC derived melanoma model^2,34,35^ (Extended Data Fig. 5e). To further test this, we performed immunofluorescence studies and were able to detect the presence of GABA itself in melanoma cells, which showed co-localization with wheat germ agglutinin (WGA), a widely used plasma membrane marker (Fig 2h). To genetically test the role of GABA-A signaling, we knocked down the GABA synthesis enzymes GAD1 and GAD2 only in melanoma cells, or the GABA-A receptor organizer ARHGEF9 (collybistin) only in keratinocytes, and then measured Cre mediated fluorescent switching (Fig 2i). While individual knockdowns showed a partial decrease in switching (Extended Data Fig. 5f and g), double knockdown of *GAD1* and *GAD2* in melanoma cells, and *ARHGEF9* in keratinocytes, showed a much larger decrease (92%, Fig 2j), highlighting the primary role of the GABAergic pathway in this form of melanoma/keratinocyte communication.

### Keratinocytes form inhibitory electrochemical synapses with melanoma cells

In the adult nervous system, GABAergic synapses are primarily involved in inhibitory neurotransmission via an influx of chloride ions into the postsynaptic cell, resulting in a decreased likelihood of a postsynaptic action potential^36,37^. Recent studies using melanocyte/keratinocyte co-cultures have demonstrated the presence of calcium spike based electrical activity between these cell types during normal development^38^. Based on this, we hypothesized that melanoma cells were reviving this developmental mechanism to promote tumor initiation by forming inhibitory GABAergic synapses with keratinocytes.

Studies in neurons have shown that synaptic GABA-A receptor activation is regulated post-transcriptionally and requires the clustering of assembly proteins like gephyrin, which ensures proper stabilization of GABA-A receptors at synapses^39^. Further, gephyrin clustering at synapses is highly dependent on the activity of collybistin (*ARHGEF9*)^30^, which we previously found was upregulated in switched keratinocytes (Extended Data Fig. 5c). To test whether melanoma/keratinocyte co-cultures expressed such synaptic markers, we performed immunofluorescence studies using gephyrin as a GABAergic postsynapse marker. We detected the presence of membrane gephyrin protein clusters specifically in keratinocytes in direct contact with melanoma cells in co-culture, highlighting the activation of the GABA-A receptor machinery in keratinocytes, only upon direct contact with melanoma cells (Figure 3a and b, Extended Data Fig. 6f). To further test this in melanoma patient samples, we stained a series of in situ melanomas in a tumor microarray to look for the presence of the GABA-A receptor machinery in the keratinocytes directly adjacent to the tumor cells. We marked melanoma cells with S100A6, and stained for gephyrin (as a marker of the GABA-A receptor), and found that n=6/6 melanoma samples contained gephyrin positive clusters in keratinocytes directly adjacent to melanoma cells as well as in more distal keratinocytes, suggesting activation of the GABAergic machinery in melanoma associated keratinocytes, but not in normal skin (Fig 3c and d). In addition, we looked at differentially upregulated pathways in switched keratinocytes using GSEA analysis. We found a strong enrichment of pathways related to synapse formation, particularly associated with the postsynapse (Extended Data Fig. 6d). We then performed electron microscopy on melanoma/keratinocyte co-cultures, which revealed striking evidence of a morphology highly consistent with synaptic structures between the two cell types, similar to what is found in neurons (Fig 3e, Extended Data Fig. 6a - c).

**Fig. 3.**
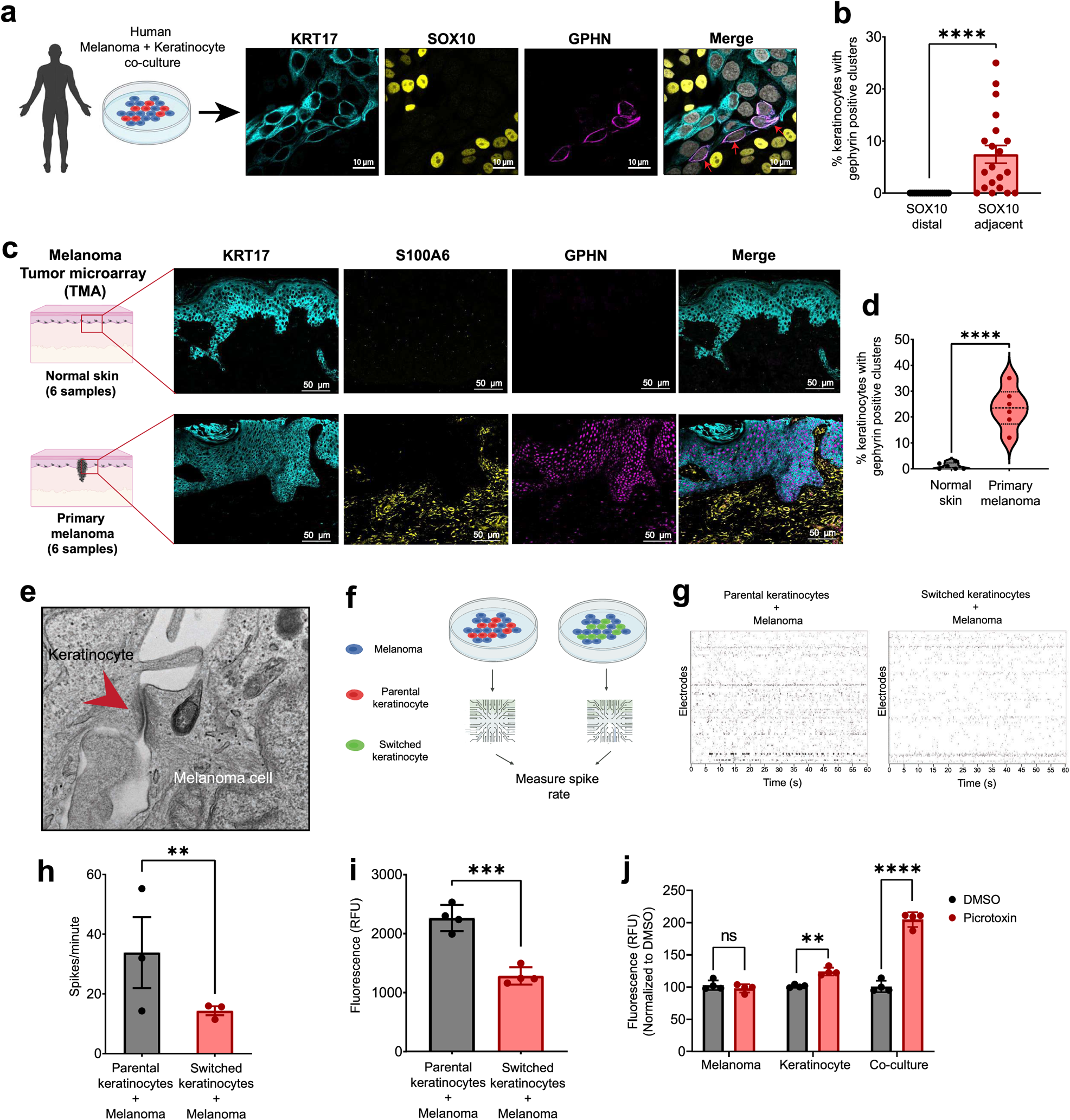
Melanoma cells form synaptic connections with keratinocytes. **(a):** Representative image of immunostaining for KRT17 (keratinocyte marker), SOX10 (melanoma marker) and gephyrin (postsynapse marker) in human melanoma/keratinocyte co-cultures. Gephyrin positive clusters are only observed in keratinocytes at sites of melanoma cell contact (indicated by red arrows). Individual cells are pseudocolored as indicated. **(b):** Quantification of gephyrin positive clusters in melanoma/keratinocyte co-cultures. Each data point represents a microscopic field quantified for the presence of keratinocyte gephyrin positive clusters pooled from 4 biological replicates (n = 20). **(c):** Representative images of a patient malignant melanoma in situ (MMIS) and normal skin sample with immunostaining for KRT17 (keratinocyte marker), SOX10 (melanoma marker) and GPHN (post-synapse marker). Individual cells are pseudocolored as indicated. **(d):** Violin plots showing % keratinocytes with gephyrin positive clusters in melanoma patient samples and normal skin. Data represent samples from n = 6 melanoma in situ patients, n = 6 normal skin, P values generated by unpaired t-test, **** p-value < 0.0001. **(e):** Transmission electron microscopy of melanoma/keratinocyte co-cultures with synapse-like structures indicated with red arrow. Representative image is shown. **(f):** Schematic for multi-electrode array (MEA) experimental setup in human melanoma co-cultures with parental or switched keratinocytes for 48 hours. **(g):** Representative spike rastergrams of 1 minute of activity on MEA chip in 48 hours melanoma/keratinocyte co-cultures. **(h):** Quantification of MEA activity in 48-hour melanoma/keratinocyte co-cultures calculated as spikes per minute. Data represent 3 biological replicates per condition with p values calculated using multiple unpaired t-test using the Holm-Šídák method for multiple comparisons, ** p-value < 0.01. **(i):** Calcium spike activity in co-cultures of melanoma cells with switched or parental keratinocytes. RFU is relative fluorescence units measured using the calcium dye, Rhod-4. Data represent 3 biological replicates per condition calculated using unpaired t-test, *** p value < 0.001. **(j):** Calcium spike activity in monocultures (melanoma and keratinocytes) and co-cultures upon picrotoxin (100 µM) addition. RFU is relative fluorescence units measured using the calcium dye, Cal-520. Data represent 3 biological replicates per condition calculated using unpaired t-test, ** p value < 0.01, *** p-value < 0.001.

To functionally test this potential synaptic relationship, we then performed extracellular electrophysiology recordings of melanoma/keratinocyte co-cultures. This allowed us to quantify fast changes in membrane voltage. We plated melanoma cells with parental keratinocytes or Cre-recombined switched keratinocytes on a multi-electrode array (MEA) system to record electrical spikes of melanoma cells alone, keratinocytes alone, or both cell types together (Fig 3f). Whereas melanoma/parental keratinocytes had substantial spiking activity (33 spikes/minute), this was greatly diminished with the melanoma/switched keratinocytes (14 spikes/minute), consistent with an inhibitory effect of GABAergic signaling present in those keratinocytes (Fig 3g and h). To further test this, we performed calcium imaging using fluorescent calcium indicators^40^, an important marker of electrical activity in neurons^41^. We loaded melanoma cells and either parental or switched keratinocytes with the calcium dye Rhod-4. Consistent with the switched keratinocytes having more GABA signaling, we found decreased calcium activity compared to parental co-cultures (35% less activity, Fig 3i). We then tested the effect of the GABA-A antagonist picrotoxin. While this had little to no effect on monocultures, we found that treatment with a GABA-A antagonist increased calcium activity in the co-cultured keratinocytes, consistent with an increase in electrical activity upon loss of inhibitory GABAergic signaling between melanoma/keratinocytes (Fig 3j).

In neurons, GABA-A receptor mediated inhibitory neurotransmission is associated with an inward flux of chloride ions, which ensures rapid hyperpolarization and decreased action potential in the postsynaptic cell. To test whether the chloride ion itself was involved in melanoma/keratinocyte communication, we used a reduced chloride medium to decrease the intracellular chloride concentration in keratinocytes, as described earlier^42^. We found that decreasing intracellular chloride ions decreased switching efficiency in co-cultures and disrupted melanoma/keratinocyte communication highlighting the critical role of the chloride influx in this form of communication (Extended Data Fig. 6e). Collectively, these data indicate that melanoma cells and keratinocytes form structures similar to neuronal synapses, and that GABAergic signaling from melanoma cells acts as an inhibitor of electrical activity in melanoma/keratinocyte co-cultures.

### GABA signaling promotes melanoma initiation

The above data suggested that melanoma/keratinocyte communication is important in melanoma initiation, and that this communication is mediated by GABA. Based on this, we wanted to genetically test the effects of GABA on melanoma initiation in vivo. To identify which GABA related proteins were most relevant, we analyzed components of the GABAergic signaling pathway in human melanoma tissue samples. We performed immunofluorescence for GAD1 (as a marker of GABA synthesis) and S100A6 (as a melanoma marker) on a human tissue microarray containing both normal skin as well as primary melanomas (n = 39 samples). Whereas nearly all normal skin samples (n = 9) were negative for GAD1, 40% of primary melanoma samples (n = 30, Fig 4a and b) were positive for GAD1. We also analyzed TCGA data, and found a remarkably strong negative correlation between high expression of GAD1 and progression free survival (Extended Data Fig. 8e). Based on this human data, we used the zebrafish to test the effect of GAD1 (and its related protein GAD2) on melanoma initiation. We used the miniCoopR system described above to activate *BRAF^V600E^* in melanocytes (along with germline p53 loss) but sensitized the system by injecting low doses of the rescue plasmids, such that most of the fish would not develop melanoma on their own (Extended Data Fig. 8a). As expected, only 10% of control fish receiving miniCoopR alone developed tumors at the 16 week time point. In contrast, transgenic fish overexpressing either gad1b (n = 53) or gad2 (n = 63) in melanocytes had a higher rate of melanoma initiation by *BRAF^V600E^* (Fig 4c and d). Furthermore, we noted that the gad overexpressing fish tended to have multiple tumors, suggesting that GAD/GABA signaling lowers the threshold for tumor initiation in this model (Extended Data Fig. 8b and c). To further test this, we then performed loss of function experiments. Because zebrafish melanomas express multiple GAD genes (*gad1a/gad1b/gad2*) ^43^ (Extended Data Fig. 7 a - d), we took advantage of the TEAZ model of transgenic melanoma in which multiplexed CRISPR cassettes can be directly electroporated into the skin of an adult fish^44^. We designed sgRNAs against all 3 zebrafish *gad* genes (*gad1a/gad1b/gad2*) and then initiated tumors using melanocyte-specific expression of *BRAF^V600E^* along with loss of *pten* and *p53* (Fig 4e). We found that CRISPR-mediated deletion of the *gad* genes resulted in a significant decrease in tumor size at both 6 weeks (Fig. 4f and g) and 10 weeks (Extended Data Fig. 8d) in the gad knockouts compared to the non-targeting control animals (65% decrease). Because GABA could be acting in a cell-autonomous manner on the melanoma cells themselves (rather than on the keratinocytes), we excluded this possibility by treating melanoma cells in vitro with GABA or picrotoxin, and found no change in proliferation rate (Extended Data Fig. 9a - b). Similarly, knockdown of GAD1/2 in melanoma cells in culture also did not affect proliferation (Extended Data Fig. 9c), indicating that the effect in vivo is not due to cell-autonomous effects on the melanoma cells themselves. Collectively, this data strongly implicates GABA signaling as a key factor in melanoma initiation in vivo via an interaction with keratinocytes.

**Fig. 4.**
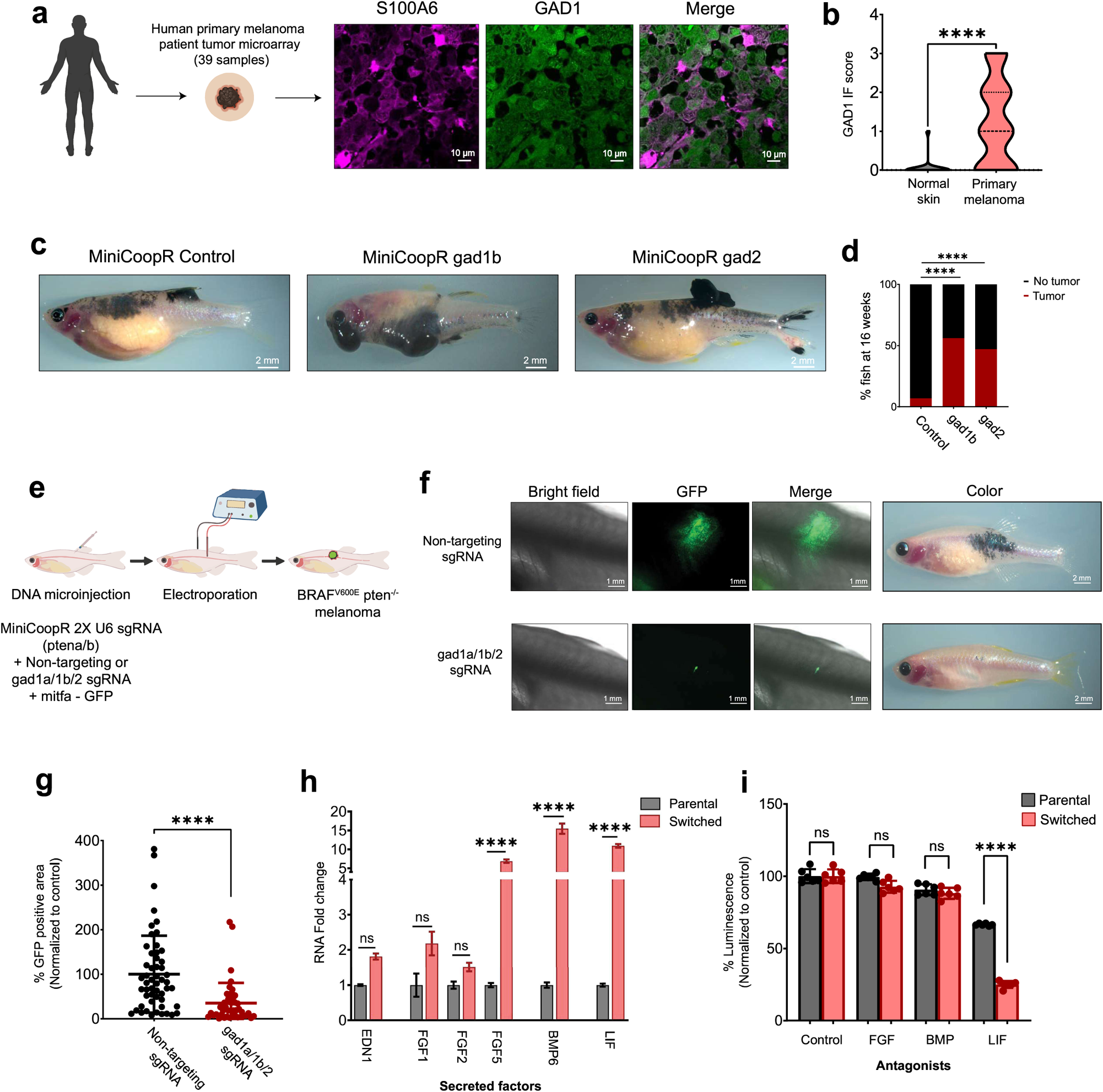
GABAergic signaling is pro-tumorigenic in melanoma. **(a):** Representative image of a patient primary melanoma sample from a primary melanoma tumor microarray (TMA) with immunostaining for S100A6 (melanoma marker) and GAD1 (GABA marker). Individual cells are pseudocolored as indicated. **(b):** Violin plots of IF (immunofluorescence) score and quantification of GAD1 immunostaining in primary melanoma tumor samples and normal skin. Data represent samples from n = 30 primary melanoma patients and n = 9 normal skin, P values generated by unpaired t-test, **** p-value < 0.0001. **(c):** Representative images of 16-week-old zebrafish with the genotype (*mitfa^−/−^ p53^−/−^ mitfa:BRAF^V600E^*) in the casper background injected with MiniCoopR rescue plasmids showing control (GFP), *gad1b* or *gad2* over-expressing tumors. **(d):** Quantification of melanoma incidence expressed as % fish with tumors in 16-week-old zebrafish over-expressing GFP or *gad1b* or *gad2* under a melanocyte specific promoter. Data represent n = 55 control (GFP) fish, n = 53 gad1b and n = 63 gad2 fish pooled from 3 biological replicates. P values generated by chi-squared test, **** p-value < 0.0001. **(e):** Schematic representation of the TEAZ based loss of function system in zebrafish to knockout GABA synthesis genes. Plasmids expressing a non-targeting sgRNA or sgRNAs against *gad1a, gad1b* and *gad2* along with MiniCoopR 2x U6 sgRNAs-pten, mitfa:Cas9 are co-electroporated to generate control (Non-Targeting) or gad KO *BRAF^V600E^ pten^−/−^* melanomas in vivo. **(f):** Representative image of a transgenic fish electroporated with melanocyte specific Cas9 and a Non-targeting sgRNA or *gad1a/gad1b/gad2* sgRNAs, 6 weeks post electroporation. **(g):** Quantification of tumor area calculated as GFP positive area and normalized to the non-targeting control plasmid group. Data represent n = 53 non-targeting sgRNA fish and n = 46 gad1a/1b/2 sgRNA fish pooled from 3 biological replicates. Error bars: SD, P values generated by Mann Whitney test, **** p-value < 0.0001. **(h):** Fold change differences in expression of secreted factors derived from RNA-seq analysis comparing switched vs parental keratinocytes. Data represent 3 biological replicates, Error bars: SD, p-values calculated using DeSeq2. **(i):** Melanoma proliferation calculated as % Luminescence in melanoma cells treated with parental or switched keratinocyte conditioned media for 48 hours +/− antagonists for the receptors of the indicated factors. Data represent 3 biological replicates (n = 12) and each condition (parental or switched) is normalized to its corresponding conditioned media control (no antagonist). Error bars: SD, p-values generated by unpaired t test with **** = p < 0.0001.

### Electrically decoupled keratinocytes promote melanoma growth through LIF

The above data suggested that the GABAergic keratinocytes were promoting the growth of the melanoma cells in vivo. Keratinocytes have previously been shown to suppress melanoma formation through physical tethering, which acts to restrain the growth of the nascent melanoma^6^. In part, this is controlled through expression of *Par3* on the keratinocytes, and its loss then allows the nascent melanoma cells to “decouple” or “escape” from the growth control of the keratinocytes^11^. In addition to physical decoupling, the keratinocytes can also express secreted factors such as *EDN1, EDN3* or *FGFs*, which can promote melanoma growth by binding to EDNRB or FGFR receptors^45,46^. Our data indicate that another way that melanoma cells could escape from keratinocyte growth control is through the activation of inhibitory GABAergic signaling, but whether these GABAergic keratinocytes promoted growth of melanoma through similar secreted factors remained unclear. To address this, we first co-cultured our human melanoma cells with parental keratinocytes or Cre-recombined switched keratinocytes and monitored the proliferation of the melanoma cells using phospho-H3 staining (Extended Data Fig. 10b). Co-culture with switched keratinocytes increased melanoma cell proliferation (Extended Data Fig. 10a and c). Further, the increase in proliferation was also seen when melanoma cells were treated only with conditioned media from the switched keratinocytes (Extended Data Fig. 10d and e), suggesting that part of the pro-tumorigenic effect could be mediated by a secreted factor from switched keratinocytes. We analyzed our RNA-seq data of the Cre-switched keratinocytes compared to the parental keratinocytes to find putative secreted ligands that would promote melanoma proliferation. While we saw no significant increase in expression of *EDN1* or *FGF1/2*, we found a significant elevation of *FGF5, BMP6* and *LIF*, factors known to promote melanoma growth^47,48,49,50^ (Fig. 4h). Based on this, we then tested whether inhibition of these pathways would slow proliferation of the melanoma cells grown with conditioned media from keratinocytes. This revealed that only inhibition of LIF receptor signaling by EC330 decreased melanoma proliferation (Fig. 4i), consistent with recent data that expression of the LIF receptor in melanoma is associated with poor prognosis in this disease^51^. Interestingly, LIF expression has been widely reported in the nervous system, including GABAergic neurons^52^, and is associated with increased growth, survival and neuroprotection^53,54^. Taken together our data suggests that loss of electrical activity between nascent melanoma cells and keratinocytes in the skin is pro-tumorigenic, and depends upon LIF mediated increase in melanoma proliferation.

## Discussion

In this study, we identify a novel mode of communication between melanoma cells and keratinocytes in the melanoma microenvironment. While these cells are closely intertwined in normal skin physiology, they typically become decoupled from each other during the early stages of melanoma formation^6,55–57^. While physical decoupling is one such mechanism, we found that keratinocytes and melanoma cells form inhibitory synapse-like structures, not previously reported in skin. Given that electrical activity is increasingly recognized to play a role in tumorigenesis^58^, it is likely that this type of GABAergic mechanism may be true in other epithelial tissues, which awaits further study.

Despite the fact that electrical activity primarily regulates cell-cell communication in excitable cells like neurons, non-excitable cells can also be regulated by electrical activity^59,60^. In the skin for example, certain ion channels regulating electrical activity and membrane potential play an important role in establishing skin pigmentation patterns^61,62^. In tumors, some of the earliest studies looking at electrical activity found that nascent tumor cells show loss of electrical activity upon transformation^63^. Further studies highlighted that the loss of electrical activity was specifically between transformed cancer cells and ‘non-transformed’ healthy cells, suggesting a functional loss of communication between tumor cells and the ‘non-transformed’ healthy cells in their microenvironment^64,65^. Our present study suggests that one of the mechanisms which could drive such loss of electrical activity-based communication in melanoma is the activation of GABAergic inhibitory signaling in the tumor microenvironment, promoting increased proliferation of nascent tumor cells via the secretion of pro-tumorigenic factors. It remains to be understood whether loss of electrical activity also invokes non-secreted mechanisms of communication.

Recent studies in brain tumors have shown that functional synapses can form between a tumor cell and a neuron which aids in tumor progression^66–68^. These studies suggest that formation of a tumor cell to neuron synapse is pro-tumorigenic both via increased neuronal activity due to stronger synaptic connections as well as via secreted paracrine factors which promote tumor proliferation. In our study, we show for the first time that functional GABAergic inhibitory synapse-like structures between skin cells can be formed in primary melanoma, independent of any input from neurons. It remains to be tested whether this unique communication pathway between melanoma cells and keratinocytes is a part of the normal physiology of melanocyte/keratinocyte communication during development. Interestingly, a recent study using GCaMP based calcium imaging in skin showed the presence of calcium spikes and ‘electric-like’ activity in melanocyte/keratinocyte co-cultures, suggesting that melanocytes and keratinocytes might communicate via electrical signals^38^. Further, a recent scRNAseq study in epidermal melanocyte populations at different stages of development found that ‘synapse formation’ is a highly upregulated pathway during melanocyte development, in addition to canonical melanocyte specific pathways like pigmentation and organelle maturation^69^. The above two studies suggest that electrical activity-based communication and ability to form synapses is closely intertwined with melanocyte development. Combined with our findings, this might indicate that nascent melanoma cells induce *GAD1* expression and use their pre-existing synaptic machinery to activate inhibitory GABAergic signaling in keratinocytes, effectively decoupling themselves from the stringent growth control of the skin keratinocytes.

An interesting finding in our study is that melanoma/keratinocyte communication is mediated by exosome-like vesicles. Our loss of function studies suggest that perturbation of exosome-like vesicle machinery in melanoma cells by pharmacological (GW4869 treatment) or genetic approaches (nSMase2 knockdown) is sufficient to block this vesicle-mediated communication between melanoma cells and keratinocytes (Extended Data Fig. 4b and c). Since most vesicle-based cargo is targeted for degradation in the recipient cell^70,71^, synaptic GABAergic signaling may be one such mechanism, which increases the magnitude of vesicle-based communication between different cell types. In accordance with this, we note that disrupting the GABA-A receptor mediated chloride efflux either by blocking the chloride channel (Fig 2c and j) or decreasing the intracellular chloride concentration (Extended Data Fig. 6e) is sufficient to disrupt this vesicle-based communication. Interestingly, drugs inducing chloride accumulation in the cell and facilitating ‘endosomal escape’ are being increasingly used for the functional delivery of extracellular vesicle cargo^72,73^. Future identification of the role of chloride in this process may pave the way for a new generation of therapeutic targets to disrupt tumor/microenvironment communication.

## Supporting information

Table S1 - LOPAC Screen Results

Table S2 - Keratinocyte RNA-seq

## Acknowledgements

The authors would like to thank members of the White lab for helpful discussions and support for this project. We thank Lee-Cohen Gould and Juan Jimenez for help with transmission electron microscopy and Chenura Jayewickreme for help with optimization of immunofluorescence protocols. We would also like to thank the MSKCC Molecular Cytology core for imaging assistance, Flow Cytometry Core for cell sorting assistance and Molecular Cytogenetics core for karyotyping assistance. All schematic figures were created using Biorender.

This work was funded with support from the NIH/NCI Cancer Center Support Grant P30 CA008748, the Melanoma Research Alliance, The Debra and Leon Black Family Foundation, NIH Research Program Grants R01CA229215 and R01CA238317, NIH Director’s New Innovator Award DP2CA186572, The Pershing Square Sohn Foundation, The Mark Foundation, The Alan and Sandra Gerry Metastasis Research Initiative at the Memorial Sloan Kettering Cancer Center, The Harry J. Lloyd Foundation, Consano and the Starr Cancer Consortium (all to R.M.W.), Melanoma Research Foundation (MRF), MSKCC TROT fellowship (both to SS), Marie-Josée Kravis Women in Science Endeavor Postdoctoral Fellowship (to MT), The Cancer Cell Map Initiative NIH / NCI U54 CA209891 (to TI).

## Author contributions

Conceptualization: MT, RMW

Methodology: MT, EH, SS, SCP, SM

Investigation: MT, EH, SS, SCP, MB, SM, TH

Visualization: MT, EH, SS, MB

Funding acquisition: RMW

Project administration: MT, RMW, TH, TI, LS

Supervision: RMW

Writing – original draft: MT, RMW

Writing – review & editing: MT, EH, SS, MB, SCP, RMW

## Data and software availability

GEO accession numbers for RNA-seq are pending. All raw data files will be made available upon request. All transgenic zebrafish lines are available upon request from the authors or via the ZIRC zebrafish stock center (https://zebrafish.org/home/guide.php). Plasmids generated in this study will be deposited to Addgene.

---

**Extended Data Fig. 1:**
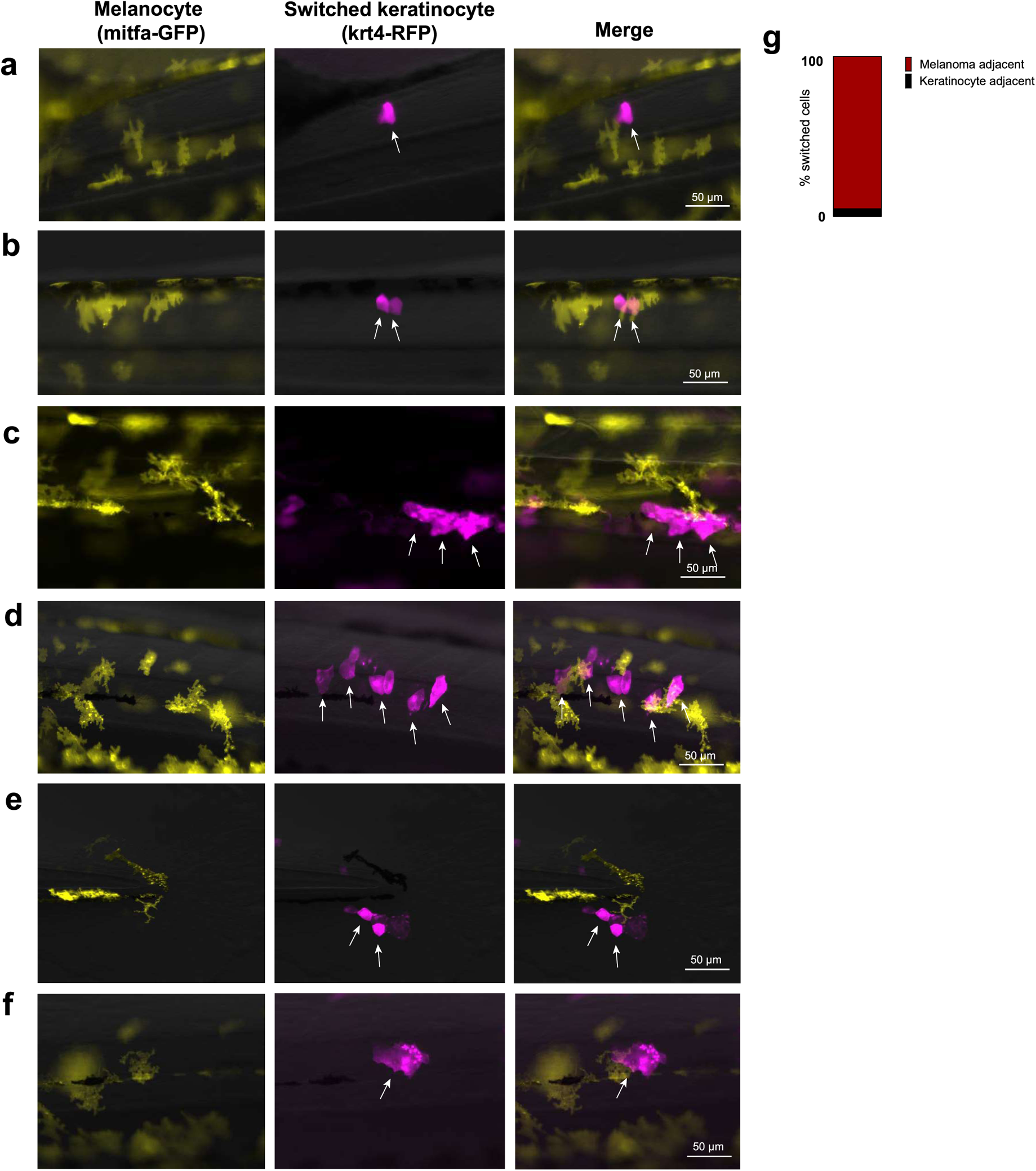
Switching requires direct contact between nascent melanoma cells and keratinocytes in vivo. **(a - f):** Representative images of 3 dpf zebrafish embryos injected with the previously described constructs (Fig. 1A). Switched keratinocytes (magenta, indicated by white arrows) are primarily located directly adjacent to a nascent melanoma cell (yellow). Individual cells are pseudocolored as indicated. **(g):** Bar plot showing % switched keratinocytes (RFP positive) in direct contact with a nascent melanoma cell (palmGFP positive). Data is pooled from 3 biological replicates (n = 60).

**Extended Data Fig. 2:**
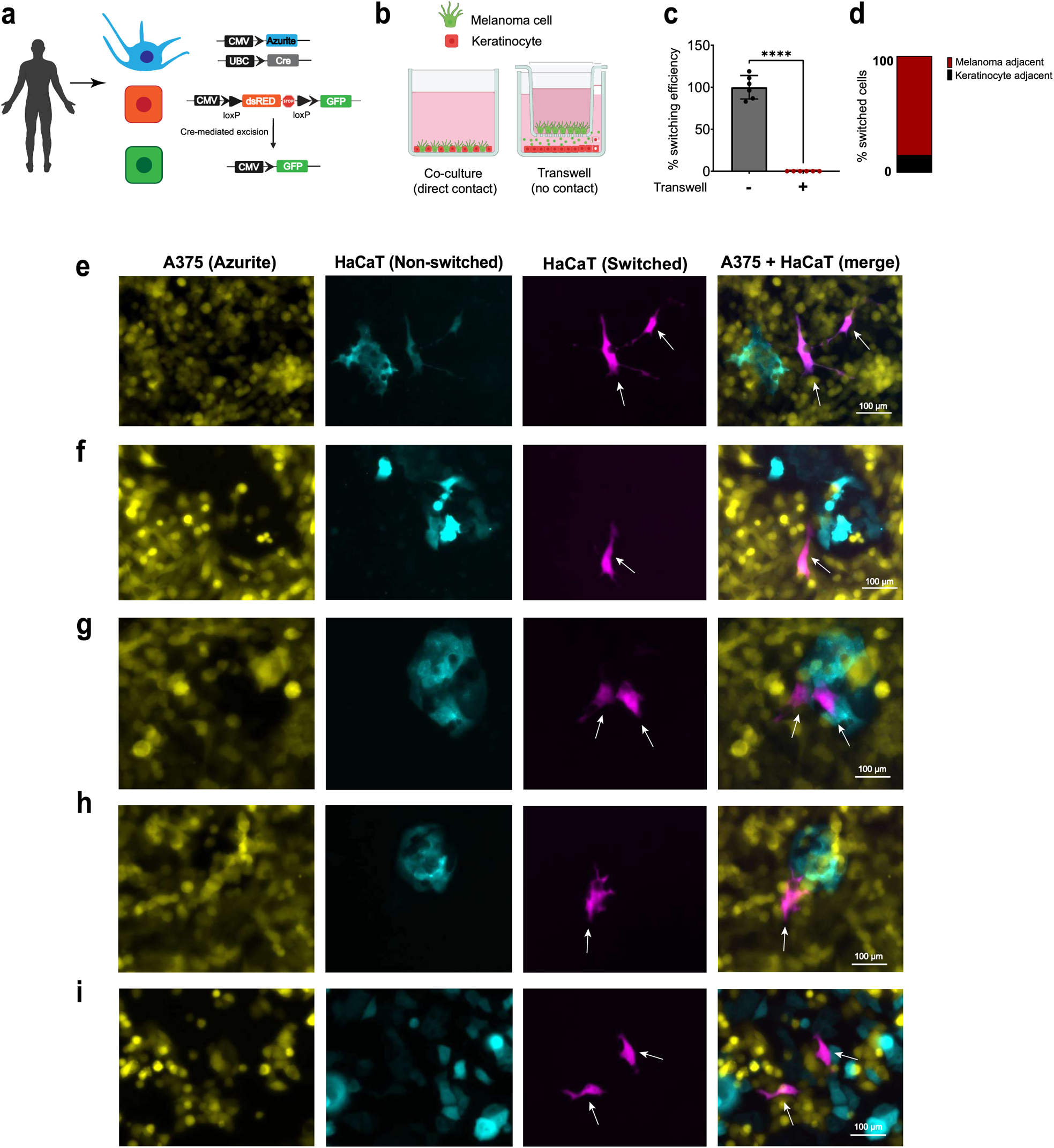
Switching requires direct contact between melanoma cells and keratinocytes in vitro. (a): Schematic representation of the genetic reporter system to detect melanoma/keratinocyte communication in human co-culture cells. (b): Schematic representation of Transwell assay for detecting melanoma/keratinocyte communication (using the Cre-loxP system) +/− direct cell-cell contact using a 400 nm transwell chamber. (c): % Switching efficiency calculated as number of switched cells per well in keratinocytes +/− direct contact with melanoma cells, normalized to no transwell (positive control). Data is pooled from 3 biological replicates (n = 6). Error bars: SD, p-values generated by unpaired t test with **** = p < 0.0001. (d): Bar plot showing % switched keratinocytes (GFP positive) in co-culture in direct contact with a melanoma cell (Azurite positive). Data is pooled from 4 biological replicates (n = 12). (e - i): Representative images of melanoma/keratinocyte co-cultures with A375 melanoma cells (yellow) over-expressing Cre and Azurite and HaCaT switched keratinocytes over-expressing the floxed dsRED to GFP switch construct (magenta, indicated by white arrows). Switched keratinocytes are primarily located directly adjacent to a melanoma cell. Individual cells are pseudocolored as indicated.

**Extended Data Fig. 3:**
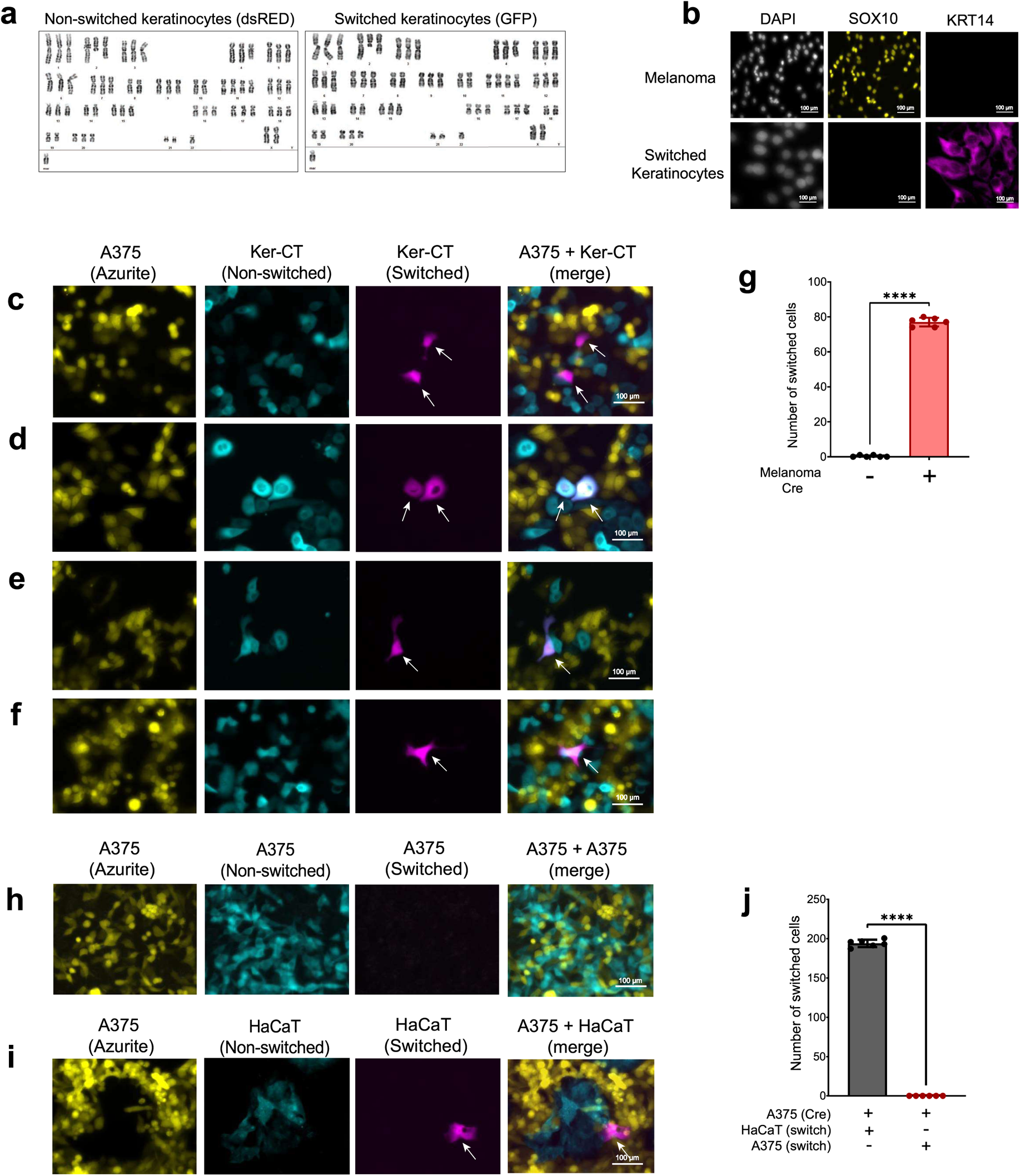
Switching does not involve cell fusion between keratinocytes and melanoma cells. **(a):** Karyotypic analysis of switched (GFP positive) vs non-switched (dsRED positive) keratinocytes post FACS sorting highlighting the presence of HaCaT specific chromosomal markers in both cell types without additional chromosomal changes in switched keratinocytes. **(b):** Immunostaining for SOX10 (melanoma marker) and KRT14 (keratinocyte marker) in monocultures of switched keratinocytes and melanoma cells to identify evidence of cell fusion between melanoma cells and keratinocytes in co-culture. Individual cells are pseudocolored as indicated. **(c - f):** Representative images of 48-hour melanoma/keratinocyte co-cultures with A375 melanoma cells over-expressing Cre and Azurite (yellow) and switched Ker-CT keratinocytes over-expressing the floxed dsRED to GFP switch construct (magenta, indicated by white arrows). Individual cells are pseudocolored as indicated. **(g):** Quantification of switching efficiency represented as number of switched cells per well in melanoma keratinocyte co-cultures with Ker-CT keratinocytes and A375 melanoma cells +/− Cre per well of a 96-well plate. Data is pooled from 3 biological replicates (n = 6). Error bars: SD, p-values generated by unpaired t test with **** = p < 0.0001. **(h - i):** Representative images of 48-hour melanoma/melanoma or melanoma/keratinocyte co-cultures with one population of A375 melanoma cells over-expressing Cre and Azurite (yellow) and a second population of A375 melanoma cells/HaCaT keratinocytes over-expressing the floxed dsRED to GFP switch construct (turquoise/magenta). Individual cells are pseudocolored as indicated. **(j):** Quantification of switching efficiency represented as number of switched cells per well in melanoma/melanoma or melanoma/keratinocyte co-cultures per well of a 96-well plate. Data is pooled from 3 biological replicates (n = 6). Error bars: SD, p-values generated by unpaired t test with **** = p < 0.0001.

**Extended Data Fig. 4:**
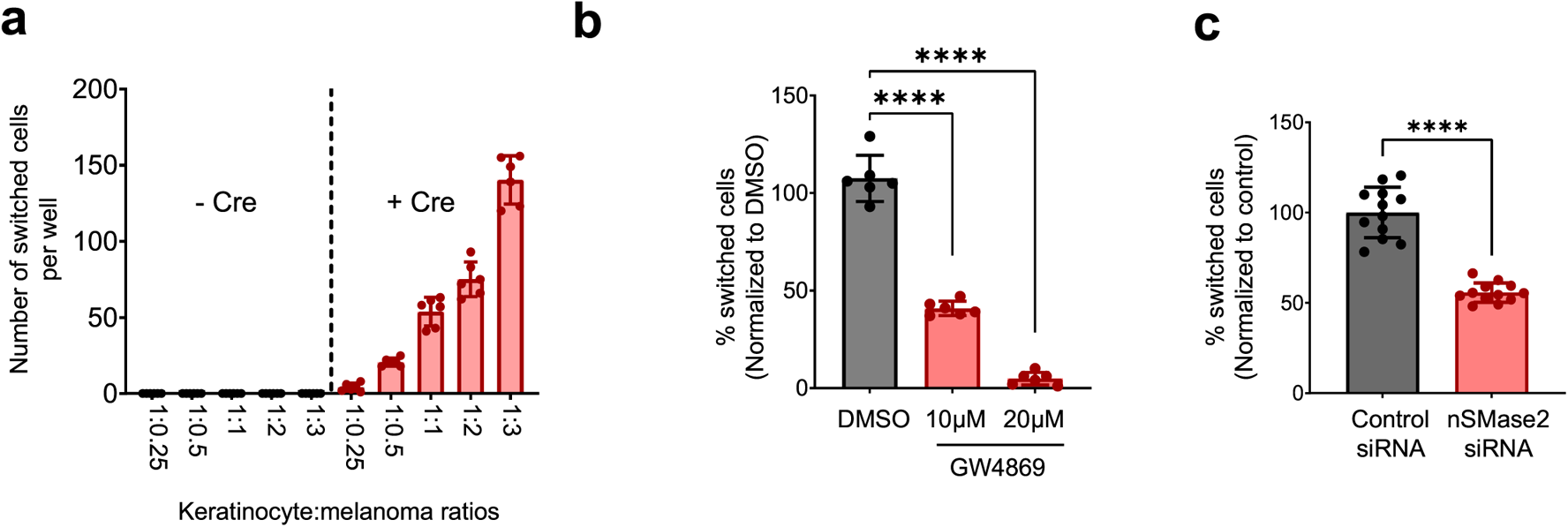
Exosome-like vesicles are transferred from melanoma cells to keratinocytes. **(a):** Switching efficiency calculated as number of switched cells per well with different melanoma/keratinocyte ratios. Increasing the number of melanoma cells, while keeping number of keratinocytes constant increases switching efficiency in keratinocytes. Data is pooled from 3 biological replicates (n = 6). Error bars: SD. **(b):** % Switching efficiency calculated as number of switched cells per well normalized to control (DMSO) upon treatment with increasing concentrations of nSMase2 inhibitor, GW4869 in melanoma/keratinocyte co-cultures for 48 hours pooled from 3 biological replicates (n = 6). Error bars: SD, p-values generated by unpaired t test with **** = p < 0.0001. **(c):** % Switching efficiency calculated as number of switched cells per well normalized to control siRNA when co-cultures are treated with nSMase2 targeting siRNA pooled from 3 biological replicates (n = 12). Error bars: SD, p-values generated by two-tailed unpaired t test with **** = p < 0.0001.

**Extended Data Fig. 5:**
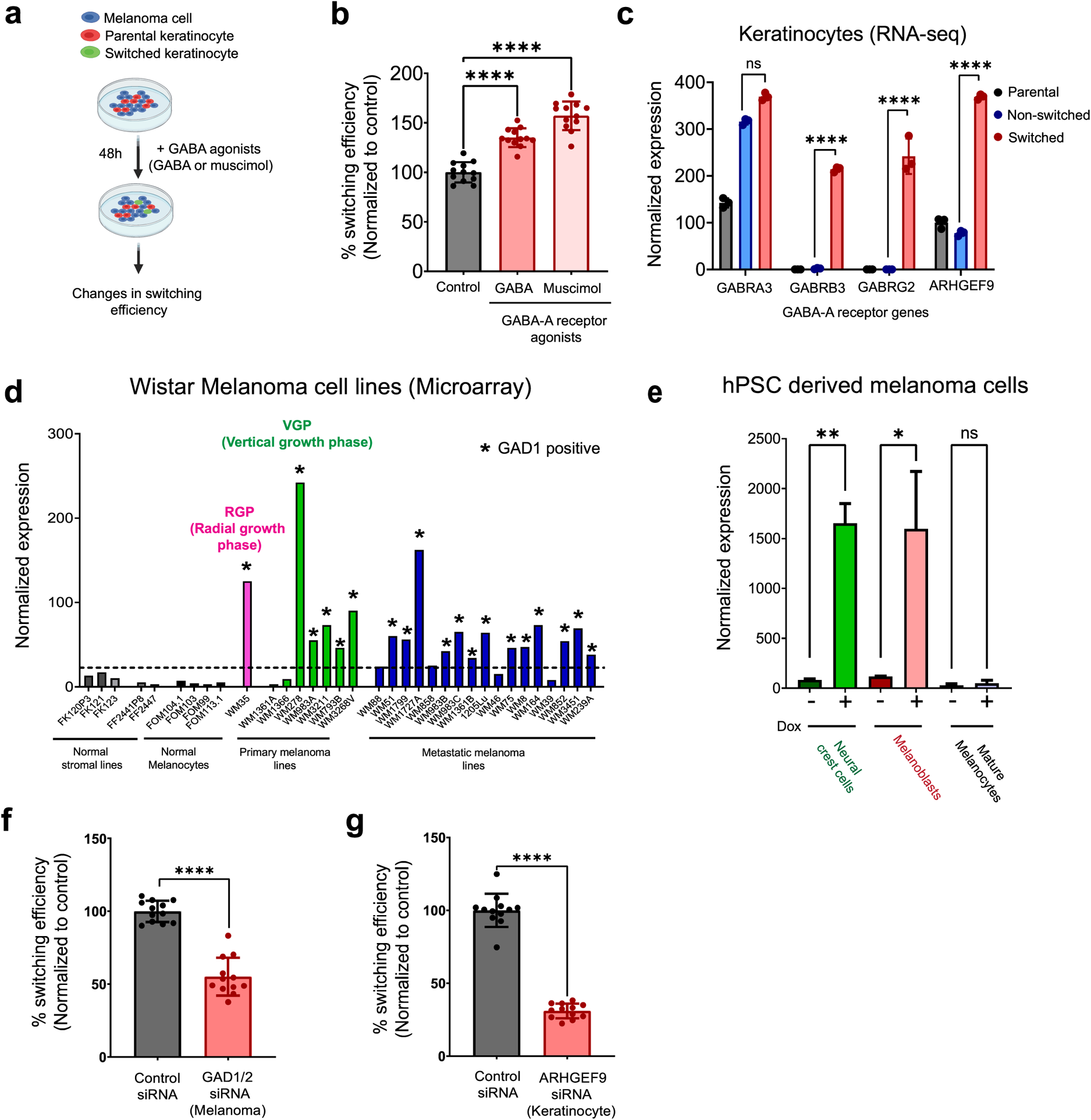
GABAergic signaling is active in skin and melanoma cells. **(a):** Schematic representation of the switching efficiency assay in vitro in melanoma/keratinocyte co-cultures grown for 48 hours +/− GABA-A agonists, GABA and muscimol. **(b):** % Switching efficiency calculated as number of switched cells per well normalized to control (DMSO) upon treatment with GABA-A agonists, GABA (100 µM) and muscimol (10 µM) in melanoma/keratinocyte co-cultures pooled from 4 biological replicates (n = 12). Error bars: SD, p-values generated by unpaired t test with **** = p < 0.0001. **(c):** Normalized expression of GABA receptor subunits (*GABRA3, GABRB3* and *GABRG2*) and collybistin (*ARHGEF9*) in parental, non-switched and switched keratinocytes from RNA-seq analysis. Data represent three biological replicates. Error bars: SD, p values were calculated using DeSeq2 with **** = p < 0.0001. **(d):** Normalized expression of *GAD1* in Wistar melanoma cell lines representing primary melanoma (RGP and VGP), metastatic melanoma, primary melanocytes (FOM lines), keratinocytes (FK lines) and fibroblasts (FF lines). * represents cell lines that are GAD1 positive. **(e):** Normalized expression of *GAD1* in triple knockout (*RB1, P53, P16*) human pluripotent stem cell (hPSC) lines differentiated into neural crest cells, melanoblasts and melanocytes (data from Baggiolini et al, 2021). Addition of doxycycline induces expression of *BRAF^V600E^* in this system. Only neural crest cells and melanoblasts induce *GAD1* expression in response to doxycycline and can form tumors in vivo. Data represent three biological replicates. Error bars: SD, p values were calculated using DeSeq2 with ** = p < 0.01 and * = p < 0.05. **(f and g):** % Switching efficiency calculated as number of switched cells per well normalized to control siRNA pooled from 3 biological replicates upon *GAD1/2* only (F) or *ARHGEF9* only (G) knockdown (n = 12). Error bars: SD, p-values generated by two-tailed unpaired t test with **** = p < 0.0001.

**Extended Data Fig. 6:**
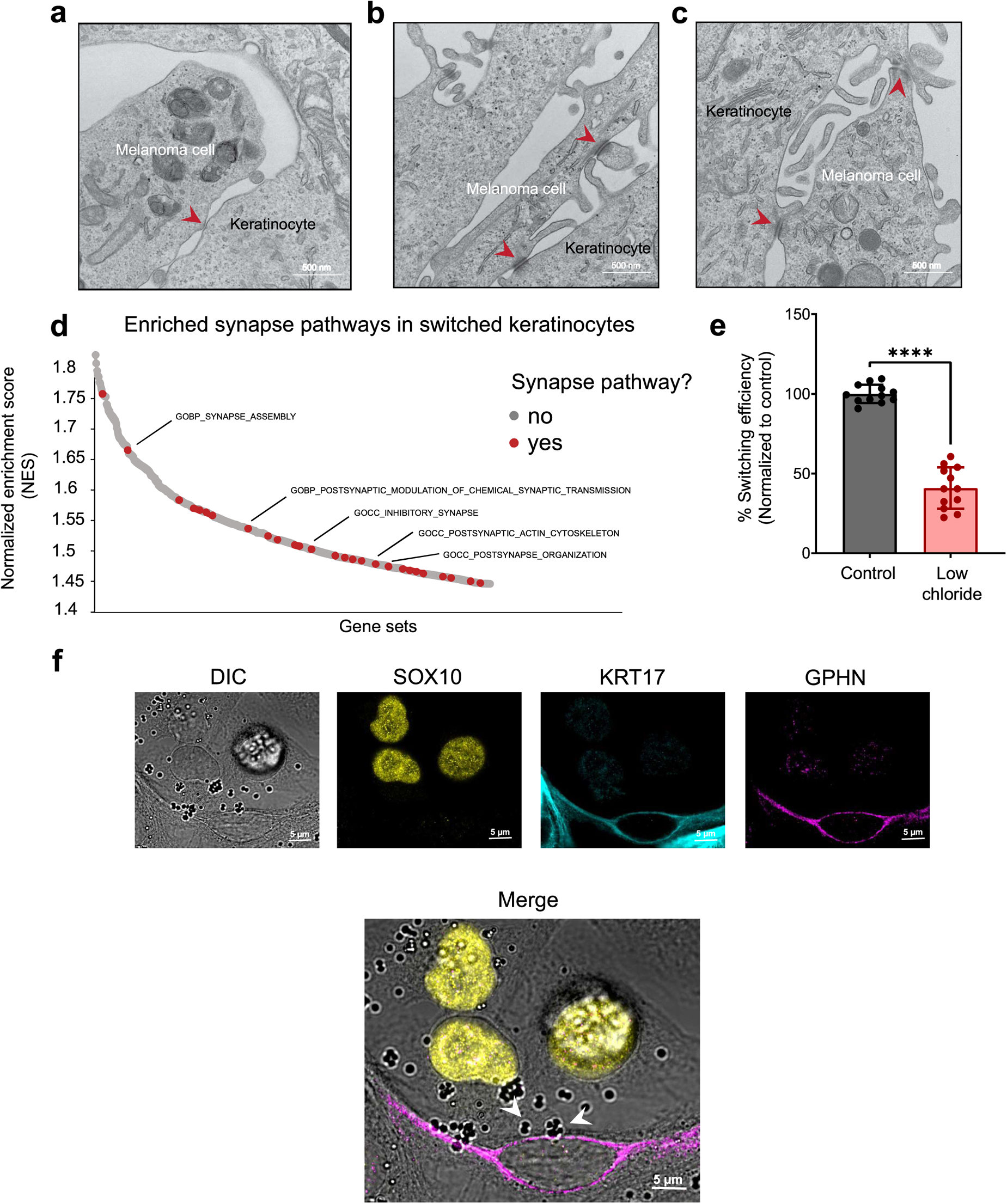
Synapse-like signatures are enriched in melanoma cells and keratinocytes. **(a - c):** Transmission electron microscopy of melanoma/keratinocyte co-cultures with synapse-like structures indicated with red arrow. Representative images are shown. **(d):** Waterfall plot of enriched pathways from GSEA analysis of switched vs parental keratinocytes. Synapse related pathways are highlighted in red. **(e):** % Switching efficiency calculated as number of switched cells per well normalized to control (regular media) upon growth in low chloride media of melanoma/keratinocyte co-cultures (to deplete intracellular chloride). Data is pooled from 4 biological replicates (n = 12). Error bars: SD, p-values generated by unpaired t test with **** = p < 0.0001. **(f):** Representative image of immunostaining for KRT17 (keratinocyte marker), SOX10 (melanoma marker) and gephyrin (post-synapse marker) in human melanoma/keratinocyte co-cultures. Individual cells are pseudocolored as indicated. DIC overlap shows gephyrin clustering in keratinocytes which are in direct contact with melanoma cells (white arrows).

**Extended Data Fig. 7:**
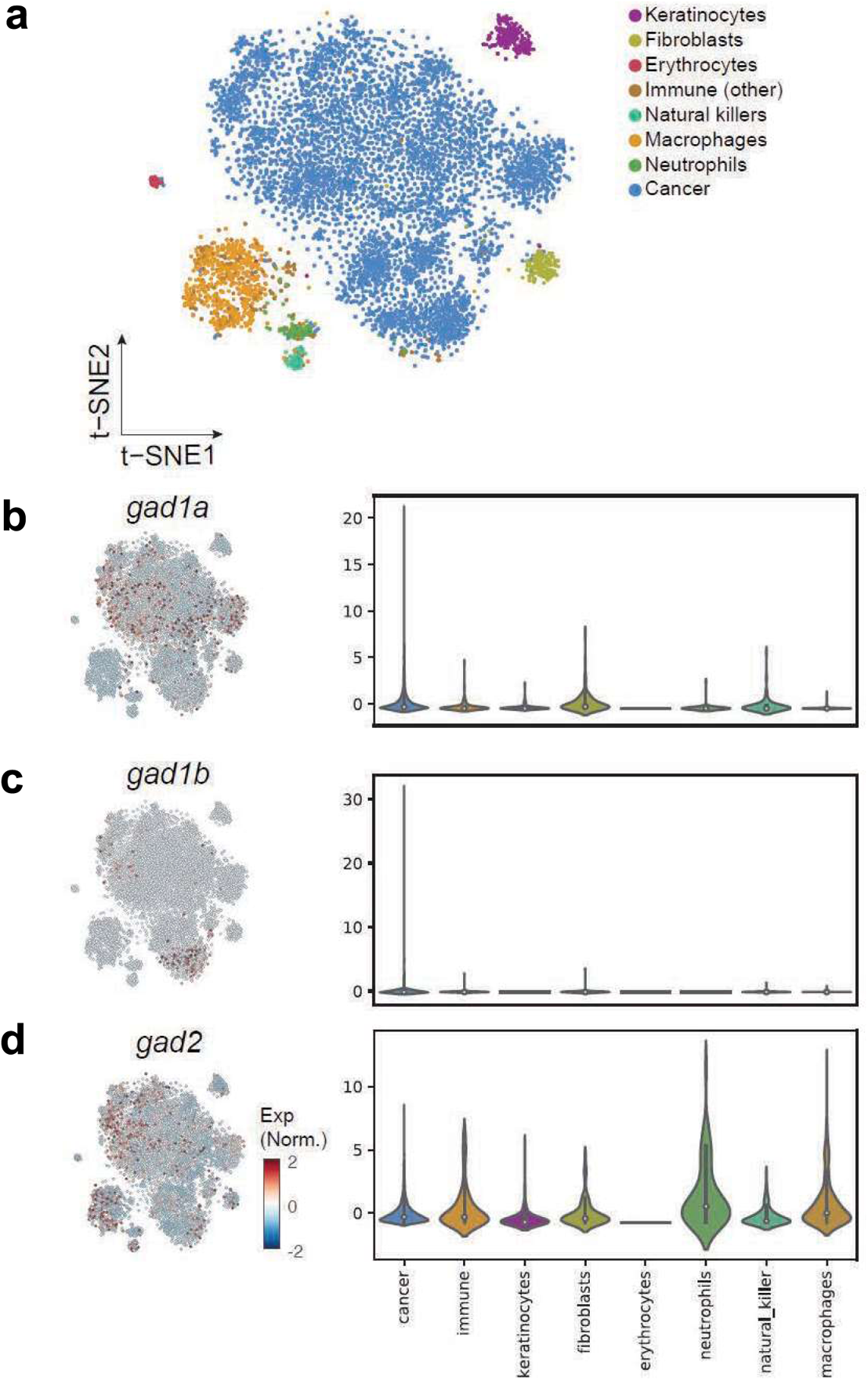
GABA producing enzymes are expressed in zebrafish melanoma cells. **(a):** tSNE analysis of 7278 individual zebrafish melanoma cells with colors indicating individual tumor and microenvironmental cell types (data from Baron et al, 2020). **(b - d):** Gene expression levels for *gad1a, gad1b* and *gad2* in single-cell RNA-seq analysis of zebrafish melanoma with individual cell clusters (left) and violin plots showing quantification of the genes in individual cell types (right). *gad1a* and *gad1b* are primarily expressed in tumor cells while *gad2* is expressed in tumor and immune cell populations.

**Extended Data Fig. 8:**
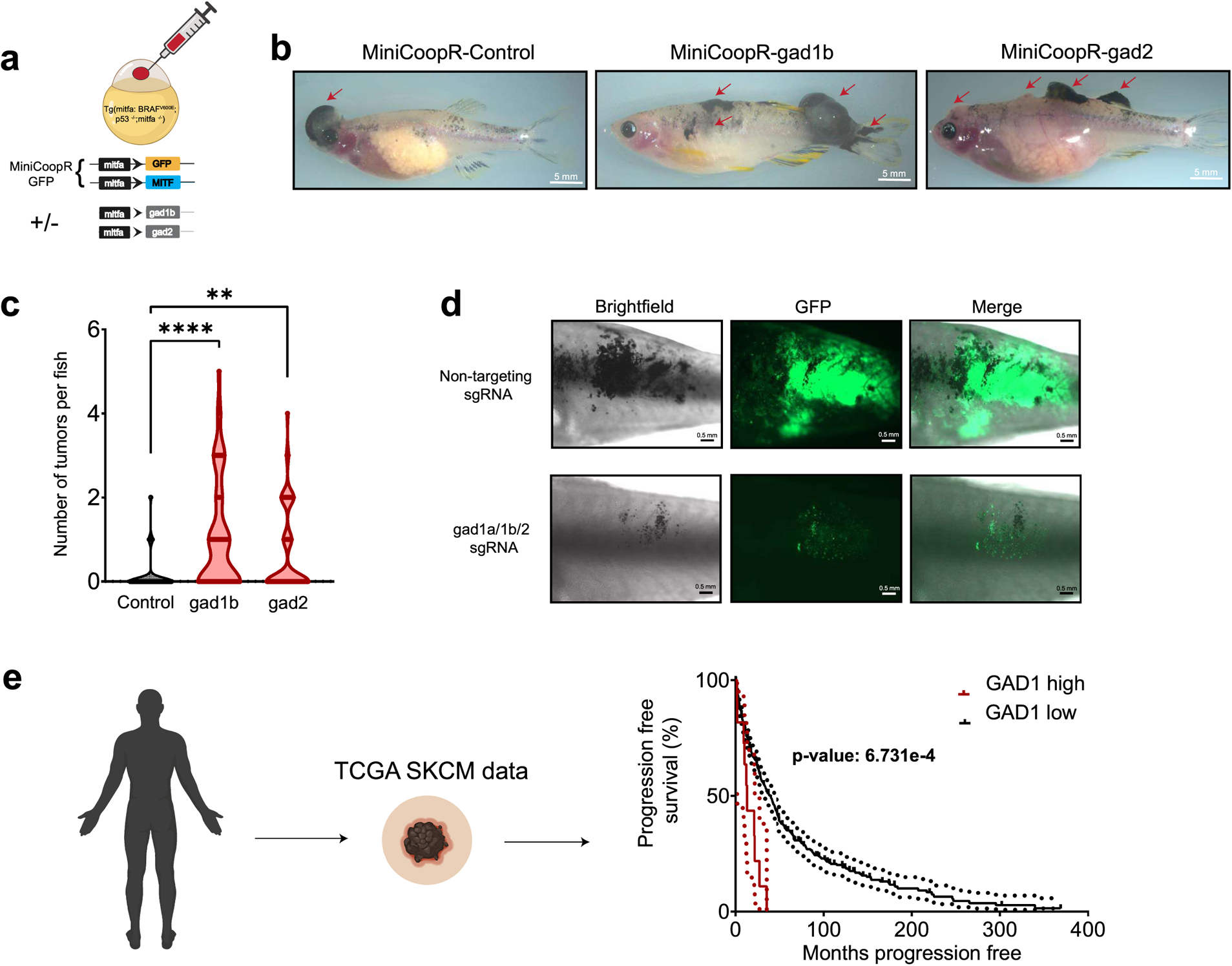
GAD expression correlates with worse prognosis in melanoma. **(a):** Schematic representation of the MiniCoopR melanocyte rescue and melanoma development system in zebrafish with over-expression of *gad1b* or *gad2* gene in a melanocyte specific manner. **(b):** Representative images of 16-week-old zebrafish injected with MiniCoopR rescue plasmids showing control (GFP), *gad1b* and *gad2* over-expressing tumors. gad over-expressing zebrafish have multiple tumors per fish (red arrow) when compared to control fish. **(c):** Violin plots showing quantification of melanoma initiation frequency expressed as number of tumors per fish in 16-week-old zebrafish over-expressing GFP or *gad1b* or *gad2* under a melanocyte specific (mitfa) promoter. Data represent n = 55 control (GFP) fish, n = 53 gad1b and n = 63 gad2 fish pooled from 3 biological replicates. P values generated by chi-squared test, ** p-value < 0.01, **** p-value < 0.0001. **(d):** Representative images of transgenic fish electroporated with melanocyte specific Cas9 and a non-targeting sgRNA or gad1a/1b/2 sgRNA, 10 weeks post electroporation. **(e):** Kaplan-Meier progression free survival curve of TCGA SKCM patients with high levels of *GAD1* expression. Log-rank p-value is indicated.

**Extended Data Fig. 9:**
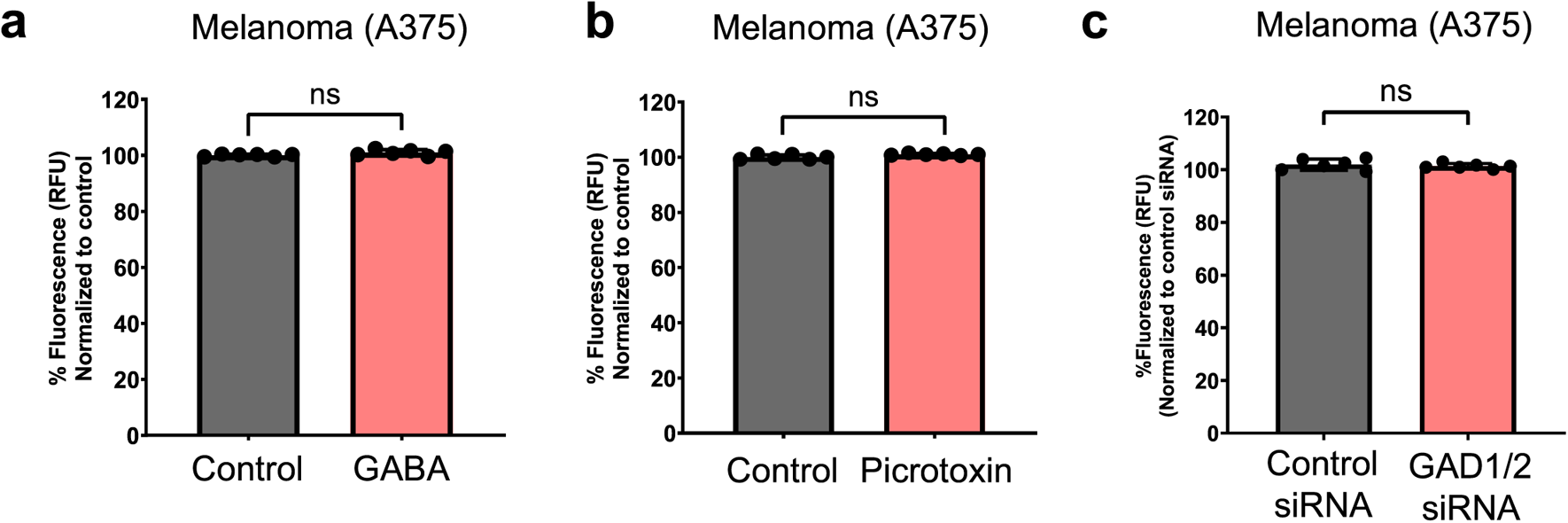
GABA treatment does not increase proliferation in monocultured melanoma cells. **(a):** Melanoma proliferation calculated as % Fluorescence in melanoma cells treated with GABA (100 µM) as indicated. Data is pooled from 3 biological replicates (n = 6) and is normalized to control treatment condition. Error bars: SD, p-values generated by unpaired t test with **** = p < 0.0001. **(b):** Melanoma proliferation calculated as % Fluorescence in melanoma cells treated with Picrotoxin (100 µM) as indicated. Data is pooled from 3 biological replicates (n = 6) and is normalized to control treatment (DMSO) condition. Error bars: SD, p-values generated by unpaired t test with **** = p < 0.0001. **(c):** Melanoma proliferation calculated as % Fluorescence in melanoma cells treated with control siRNA or GAD1/2 siRNA as indicated. Data is pooled from 3 biological replicates (n = 6) and is normalized to control siRNA treatment condition. Error bars: SD, p-values generated by unpaired t test with **** = p < 0.0001.

**Extended Data Fig. 10:**
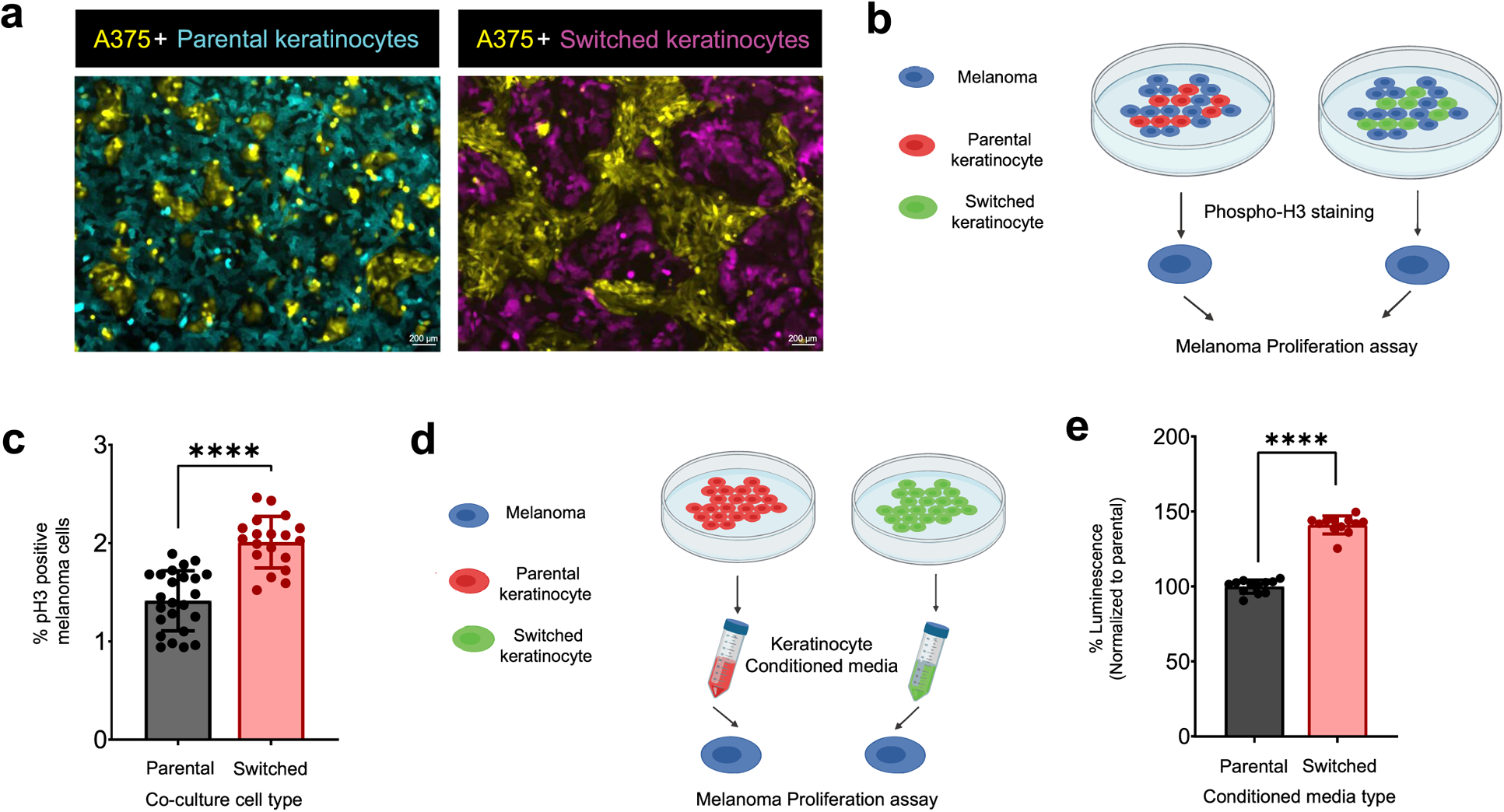
Switched keratinocytes are pro-tumorigenic in melanoma. **(a):** Representative images of melanoma/keratinocyte co-cultures with A375 melanoma cells (yellow) and HaCaT parental keratinocytes (turquoise, left) or HaCaT switched keratinocytes (magenta, right). Switched keratinocytes are pro-tumorigenic in melanoma as seen by the increased number of A375 cells in co-culture with switched keratinocytes (right panel). **(b):** Schematic representation of melanoma proliferation assay in melanoma/keratinocyte co-cultures using phospho-histone (pH3) staining. **(c):** Melanoma proliferation calculated as % phospho-histone (pH3) positive melanoma cells when co-cultured with parental or switched keratinocytes for 48 hours. Data is pooled from 4 biological replicates (n = 12). Error bars: SD, p-values generated by unpaired t test with **** = p < 0.0001. **(d):** Schematic representation of luminescence-based melanoma proliferation assay. Melanoma cells are treated with 24-hour conditioned media from parental or switched keratinocytes for 48 hours and proliferation is measured using the CellTiter-Glo assay. **(e):** Melanoma proliferation calculated as % Luminescence in melanoma cells treated with conditioned media as indicated. Data is pooled from 4 biological replicates (n = 12) and is normalized to parental keratinocyte condition. Error bars: SD, p-values generated by unpaired t test with **** = p < 0.0001.

**Extended Data Fig. 11:**
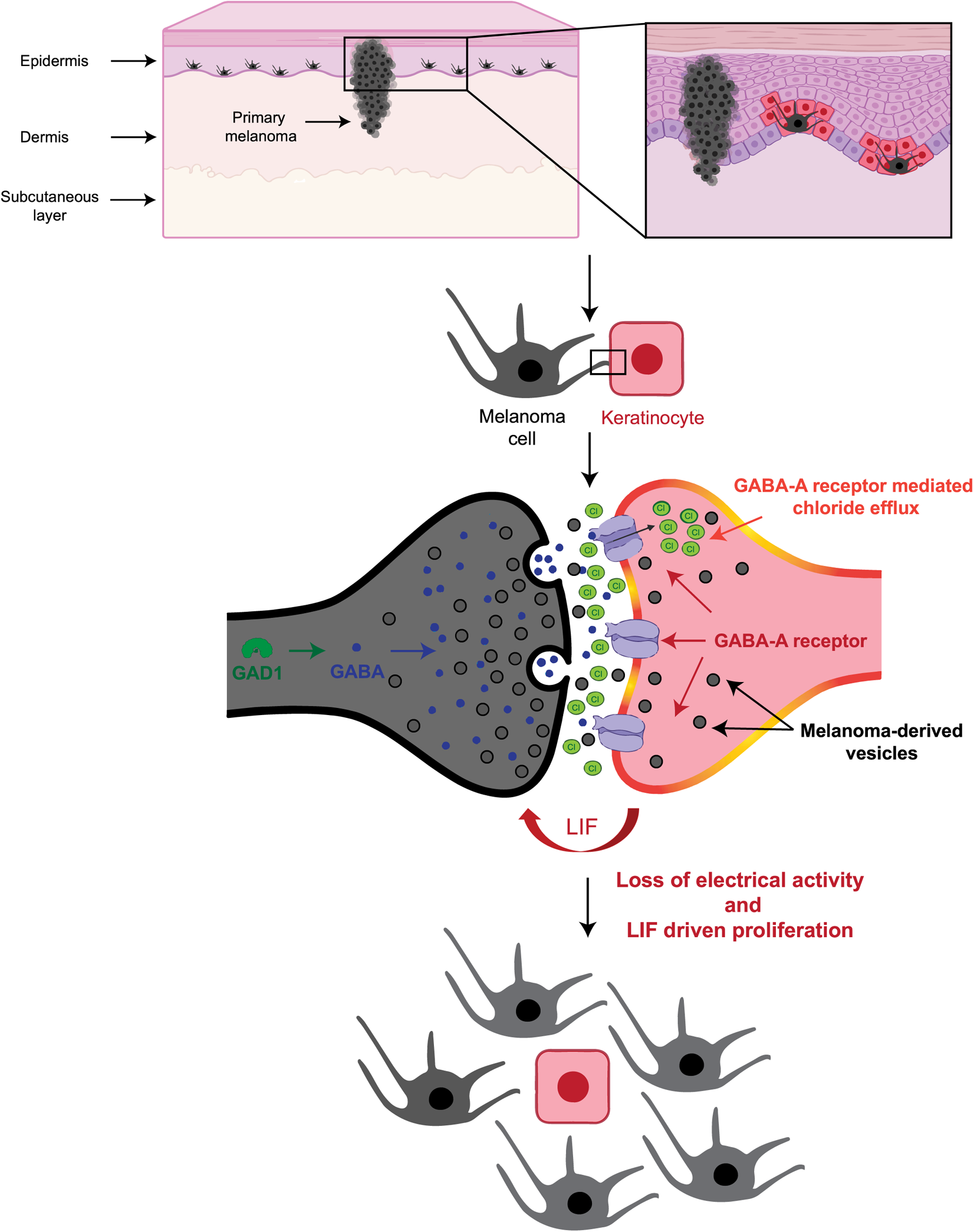
Model for GABAergic signaling driven melanoma/ keratinocyte communication.

## Methods

### Zebrafish

#### Zebrafish husbandry

All zebrafish experiments were carried out in accordance with institutional animal protocols from Memorial Sloan Kettering Cancer Center (MSKCC) Institutional Animal Care and Use Committee (IACUC), protocol number 12-05-008. Fish stocks were kept under standard conditions at 28.5 under 14 hr light/10 hr dark cycles, with salinity and pH (7.4) controlled conditions. Animals were fed standard zebrafish diet consisting of brine shrimp followed by Zeigler pellets. Embryos were collected from natural mating and incubated in E3 buffer (5 mM NaCl, 0.17 mM KCl, 0.33 mM CaCl_2_, 0.33 mM MgSO_4_) at 28.5. All anesthesia was performed using Tricaine-S (MS-222, Syndel USA, 712 Ferndale, WA) with a 4g/L, pH 7.0 stock. Sex determination in embryos is not possible at 3-5 days post fertilization (dpf).

#### In vivo switch assay in zebrafish embryos

For the in vivo switch assay in zebrafish, one cell stage casper ^74^ mitfa:BRAF^V600E^ p53^−/−^ embryos (30 embryos per condition) were injected with the following reporter constructs:

1. krt4-loxP-lacZ-loxP-tdTomato OR krt4-loxP-GFP-loxP-tdTomato
2. MiniCoopR-palmGFP OR MiniCoopR-empty
3. mitfa-cre OR mitfa-empty

at 5ng/µl with tol2 mRNA at 20ng/µl. Embryos were screened for GFP fluorescence and imaged at 3 dpf to measure GFP and tdTomato (RFP) fluorescence. To calculate switching efficiency in embryos, we used GFP fluorescence from krt4-loxP-GFP-loxP-tdTomato construct to mark all rescued keratinocytes in the embryo and tdTomato (RFP) fluorescence to mark switched keratinocytes only. For quantifying switching efficiency, background autofluorescence was subtracted and rescued keratinocyte area was quantified by uniformly thresholding the GFP intensity using the default ImageJ segmentation algorithm, followed by quantifying the switched keratinocyte area by uniformly thresholding the tdTomato (RFP) intensity across all animals. % Switching efficiency was calculated as tdTomato (RFP) positive area/GFP positive area times 100 using ImageJ (NIH). For cell-cell direct contact measurements in Fig S1A, each embryo was manually screened for the presence of switched keratinocytes (tdTomato positive from the construct krt4-loxP-lacZ-loxP-tdTomato) adjacent or not adjacent to a rescued melanocyte (GFP positive from the construct MiniCoopR-palmGFP). All images were acquired on a Zeiss AxioZoom V16 fluorescence microscope.

#### Pharmacological treatment of zebrafish embryos

Zebrafish melanoma prone embryos (casper mitfa:BRAF^V600E^ p53^−/−^) were injected with the following constructs: krt4-loxP-GFP-loxP-tdTomato + MiniCoopR-Cre (20 embryos per condition) at the one-cell stage. At 8 hours post fertilization (hpf) embryos were placed in a 40 µm cell strainer in a 6-well dish in 6 ml of E3 water. Each well was treated with the following compounds: DMSO or Muscimol (Sigma #M1523) at 10 µM, or Picrotoxin (Sigma #P1675) at 100 µM. The compounds were added at 8 hpf and reapplied at 1 dpf and 2 dpf. Embryos were imaged at 3 dpf using the same imaging and quantification protocol as described above to calculate % Switching efficiency.

#### Generation of gad1b and gad2 overexpressing MiniCoopR transgenic fish

Transgenic zebrafish were generated by injection of 5 ng/µl of MiniCoopR-GFP and empty vector or MiniCoopR-GFP; mitfa-gad1b or MiniCoopR-GFP; mitfa-gad2 and tol2 mRNA at 20 ng/µl in melanoma prone casper mitfa:BRAF^V600E^ p53^−/−^ embryos. Embryos were screened for melanocyte rescue at 5 dpf. Embryos with successful melanocyte rescue were grown to adulthood and scored for the emergence of tumor at 10 and 16 weeks post fertilization (wpf).

#### TEAZ electroporation

Transgene Electroporation in Adult Zebrafish (TEAZ) was utilized to generate melanomas as previously described^44^. casper mitfa:BRAF^V600E^ p53^−/−^ transgenic animals (5-6 months old) were electroporated with plasmids to generate BRAF^V600E^, p53^−/−^, pten^−/−^ tumors expressing either a Non-targeting sgRNA or sgRNAs against gad1a, gad1b and gad2. Plasmids electroporated per animal included MiniCoopR 2x U6 sgRNAs-pten, mitfa:Cas9 (300 ng)^75^ to induce melanocyte rescue and loss of pten, mitfa:GFP (125 ng), Tol2 (58 ng) and 100 ng of Non-targeting or gad sgRNAs. Briefly, adult male fish were anesthetized with 0.2% tricaine and injected with 1µl of plasmid mix described above into the skin below the dorsal fin. Fish were electroporated and allowed to recover in fresh system water. Electroporation was performed using a CM830 Electro Square Porator from BTX Harvard Apparatus and Genepaddles 3×5mm, with a voltage of 45V, 5 pulses, 60ms pulse length and 1s pulse interval. Fish were imaged every week using brightfield and fluorescent imaging at 25X and 10X using a Zeiss AxioZoom V16 fluorescence microscope. Tumor area was quantified by GFP fluorescence using ImageJ.

All gad sgRNAs for this experiment were designed using CHOPCHOP^76^ and GuideScan^77^. The sgRNA sequences are outlined below:

Non-targeting: AACCTACGGGCTACGATACG

ptena: GAATAAGCGGAGGTACCAGG

ptenb: GAGACAGTGCCTATGTTCAG

gad1a: TGACGTCACCTATGACACGG

gad1b: TACGACAACCTGCCACAAGT

gad2: GTAGAGATCCGAAAAGCACG

#### Plasmid construction

For melanocyte-specific constructs, the previously developed mitfa promoter^78^ in a gateway compatible 5’ entry vector was used. For keratinocyte-specific constructs, we used the previously described krt4 promoter which labels differentiated keratinocytes in the zebrafish skin^79^. The keratinocyte-specific switch construct was generated by modifying the previously generated ubiquitin switch construct in zebrafish, ubb-loxP-GFP-loxP-tdTomato^80^. The following plasmids were constructed using the Gateway Tol2 kit^81^ :

1. krt4:loxP-lacZ-loxP-tdTomato-394
2. krt4:loxP-GFP-loxP-tdTomato-394
3. krt4:loxP-GFP-loxP-DTA-394
4. MiniCoopR:palmGFP
5. MiniCoopR: Cre
6. mitfa:palmGFP-394
7. mitfa:cre-394
8. ubb:cre-394bsd (blasticidin in 394 backbone)
9. mitfa:gad1b-394
10. mitfa:gad2-394

For cloning gad1a, gad1b and gad2 targeting sgRNAs into a plasmid backbone, we used three different U6 promoters as described in^82^ and cloned all the sgRNAs (Non-targeting or gad) in a single gateway compatible 394 backbone with tol2 arms. Guide RNA cutting efficiency was validated using the Surveyor mutation kit (IDT #706020). MiniCoopR 2xU6:gRNA, mitfa:Cas9 (MAZERATI)^75^ was a gift from Leonard Zon (Addgene plasmid #118844).

### Cell culture

#### Melanoma cell culture

Human melanoma cell lines (A375, A375-MA2, HS294T, A101D, SKMEL24, SKMEL3, SKMEL5, C32, SH-4) were obtained from ATCC where routine cell line authentication is performed. The zebrafish melanoma cell line (ZMEL1) was derived from a tumor in a mitfa-BRAF^V600E^ p53^−/−^ zebrafish as described previously^83^. Mouse melanoma cell lines, YUMM 1.1 and YUMM 4.1^84^ were a kind gift from the Neal Rosen lab. Human melanoma cell lines, A375, A375-MA2, Hs294T, A101D and SH-4 were maintained in DMEM (Gibco #11965) supplemented with 10% FBS (Gemini Bio), 1X penicillin/streptomycin (Gibco #15140122), SKMEL-24 and C32 were maintained in EMEM (ATCC #30-2003) supplemented with 10% FBS (Gemini Bio), 1X penicillin/streptomycin (Gibco #15140122), SKMEL-3 was maintained in McCoy’s 5a Medium Modified (ATCC #30-2007) supplemented with 10% FBS (Gemini Bio), 1X penicillin/streptomycin (Gibco #15140122). Mouse melanoma lines, YUMM 1.1 and YUMM 4.1 were maintained in DMEM-F12 (ATCC #30-2006) supplemented with 10% FBS (Gemini Bio), 10% Non-essential Amino Acids (Gibco #11440-076), 1X penicillin/streptomycin (Gibco #15140122). Zebrafish melanoma cell line, ZMEL1 was maintained in DMEM (Gibco #11965) supplemented with 10% FBS (Gemini Bio), 1X penicillin/streptomycin/glutamine (Gibco #10378016), and 1X GlutaMAX (Gibco #35050061) and grown at 28°C with 5% CO_2_ in a humidified incubator. All cells were passaged less than 20 times before a low passage batch was thawed. They were routinely tested for mycoplasma using a luminescence-based mycoplasma detection kit (MycoAlert Mycoplasma Detection kit, Lonza #: LT07-318).

#### Keratinocyte cell culture

HaCaT keratinocye cell line^85^ was obtained from Addexbio and authenticated at the MSKCC Molecular Cytogenetics Core. Ker-CT is an hTERT immortalized keratinocyte cell line^86^ and was obtained from ATCC. HaCaT cells were maintained in DMEM (Gibco #11965) supplemented with 10% FBS (Gemini Bio), 1X penicillin/streptomycin (Gibco #15140122). Ker-CT cells were maintained in KGM-Gold Keratinocyte growth medium supplemented with KGM-Gold™ BulletKit™(Lonza #00192060). Cells were split when confluent, approximately 2X per week, and were used directly in co-cultures. For melanoma/keratinocyte co-cultures, 50:50 media from melanoma cells and keratinocytes was used and co-cultures were maintained for 48 hours to three weeks. For low chloride media experiments, Na-gluconate and K-gluconate was used as a substitute for chloride as described in^42^ to maintain isotonicity.

#### Cell line generation

All human and mouse melanoma lines were engineered to overexpress Cre under the UBC promoter modified from Addgene plasmid #65727 as described in^87^. Zebrafish ZMEL1-Cre overexpressing lines were generated using electroporation with the plasmid ubb:Cre-394Bsd using the Neon transfection system (Thermo Fisher). All human and mouse melanoma lines were selected with puromycin (1µg/ml) while the zebrafish ZMEL1 line was selected with blasticidin (4µg/ml) for three weeks. For switch cell line generation in melanoma cells (A375) and keratinocytes (HaCaT and Ker-CT), the plasmid, pLV-CMV-LoxP-DsRed-LoxP-eGFP (Addgene plasmid #65726) was used for lentiviral infection as described in^87^. All switch lines were subjected to puromycin selection followed by two rounds of FACS sorting to eliminate any double positive (dsRED/GFP positive) cells.

#### In vitro switch assay in melanoma/keratinocyte co-cultures

HaCaT or Ker-CT reporter keratinocytes expressing the switch cassette (0.4 million cells) and all human/mouse/zebrafish melanoma cell lines expressing Cre (1.2 million cells) were seeded in 1:3 ratios in 6-well plates with 50:50 keratinocyte/melanoma cell media. For zebrafish melanoma (ZMEL1) and human keratinocyte (HaCaT) co-cultures, cells were maintained in a 28 humidified incubator with 5% CO_2_. For all other co-cultures, cells were maintained in a 37 with 5% CO_2_ humidified incubator. After 48 hours of co-culture, conditions were blinded and each well was manually scored for the number of switched keratinocytes (GFP positive) per condition. Control co-cultures with no Cre melanoma cells were scored in parallel to eliminate the possibility of background switching in each condition. All imaging was performed in a Zeiss AxioObserver fluorescence microscope.

#### siRNA treatment

For siRNA studies, Dharmacon ON-TARGETplus SMARTpool siRNAs were used to knockdown individual human genes and cells were treated based on the manufacturer’s instructions. Briefly, melanoma cells or keratinocytes were seeded overnight in regular media with no antibiotics in 6-well dishes as monocultures and transfected with the indicated SMARTpool siRNAs and DharmaFECT 1 transfection reagent (Horizon #T-2005-01) in serum free media. 72 hours post-transfection, siRNA treated cells (melanoma cells or keratinocytes) were treated with a second dose of siRNA, followed by a 48 hour incubation and then co-cultured with the corresponding non-targeting siRNA treated melanoma cells or keratinocytes and incubated for an additional 48 hours. Knockdown efficiency of monocultures was measured at 72 hours post transfection. For calculating switching efficiency, number of switched cells per condition (3 technical replicates) was counted 48 hours post co-culture and 7 days post transfection. Knockdown efficiency was validated using qPCR at 5 days and Western Blot at 7 days post transfection. All siRNA-treated cells were monitored for signs of toxicity or changes in proliferation rate using Cyquant Cell Proliferation Assay (see below) and no toxicity was observed for the indicated siRNA treatments.

#### Transwell assay

Transwell assay was performed to detect requirement of cell-cell contact for switching in keratinocytes, as described in^87^. Briefly, 20,000 HaCaT keratinocytes expressing the switch construct were plated alone or in combination with 60,000 A375 melanoma cells expressing Cre per well of a 24-well plate. 24 hours post plating, 20,000 A375 melanoma cells (+/− Cre) were seeded into the upper Transwell chamber (400 nm size) with a layer of keratinocytes seeded previously at the bottom. 48 hours post seeding the transwell chamber, switched GFP positive cells were counted per condition. Cells were incubated for an additional two weeks and scored for GFP positive switched keratinocytes in the bottom well.

#### Sigma LOPAC library small molecule screen

For the small molecule screen, we wanted to identify enhancers of switching efficiency, which is a readout of melanoma/keratinocyte communication, in co-cultures, using the in vitro switch assay outlined above. To do this, human melanoma/keratinocyte co-cultures (HaCaT and A375 cells) were seeded in 96-well plates at 1:3 ratios on Day 0. 24 hours post seeding, co-cultures were treated with 10 µM chemicals from the Sigma LOPAC 1280 library. All chemicals were prepared from 10 mM stocks and diluted in cell growth medium. Eight DMSO only wells were included in each plate to quantify basal switching efficiency. Co-cultures were incubated for an additional 48 hours post chemical addition and switching efficiency was quantified 3 days post seeding and 48 hours post adding chemicals. Each well was manually screened and the number of GFP positive switched keratinocytes was counted. Fold change was calculated by dividing the number of switched keratinocytes per well to the average number of switched keratinocytes per well in DMSO controls. After the identification of positive hits, a majority of which were modulators of the GABA-A signaling pathway, a second round of validation was performed in a 6-well format, using bonafide GABA-A receptor agonists like GABA (Sigma #A2129), Muscimol (Sigma #M1523), as well as GABA-A receptor antagonists like Bicuculline methbromide (Sigma #B7561) and Picrotoxin (Sigma #P1675), followed by in vivo validation in zebrafish embryos.

### RNA-seq of keratinocytes

HaCaT keratinocyte cell line expressing the switch construct was grown either in monoculture or co-culture with A375 melanoma cells expressing Cre in a 1:3 keratinocyte/melanoma ratio for 21 days in complete DMEM. Co-cultures were split 1X per week upon reaching confluence. Monoculture keratinocytes (dsRED positive) and co-culture keratinocytes, both switched and non-switched (GFP positive and dsRED positive) were then isolated using FACS and plated on 6-well dishes for recovery following the FACS procedure. Total RNA was isolated from the FACS isolated keratinocytes post-recovery using the Quick RNA miniprep kit (Zymo) and purified RNA was delivered to GENEWIZ (South Plainfield, NJ) for mRNA preparation with the TruSeq RNA V2 kit (Illumina) and 150bp paired-end sequencing on the Illumina HiSeq2500. Quality control of the raw reads from RNA-seq fastq files was performed using FASTQC (Babraham Bioinformatics) and trimming was performed using TRIMMOMATIC^88^. Trimmed reads were mapped to the human (hg38) genome using STAR^89^. Gene counts of aligned reads were performed using the feature counts algorithm^90^, followed by differential gene expression analysis using DeSeq2^91^. Pathway and gene ontology analysis was performed using GSEA^92^. All data will be deposited in the GEO database.

### Immunostaining

#### Immunofluorescence staining of cultured cells

HaCaT/A375 co-cultures were grown on 35 mm glass bottom dishes (ThermoFisher #150682) for 48 hours. Post 48 hours incubation, growth media was aspirated from the dishes and cells were fixed with 4% PFA for 20 min at room temperature (RT) and washed 3X with PBS. Following this, cells were permeabilized for 15 min in 0.1% Triton X-100 (diluted from 10% stock, Sigma #93443) in 1X PBS, followed by 3X washes with PBS. Blocking was performed using 10% normal goat serum (ThermoFisher #50062Z) for 1 hour at RT followed by overnight incubation with primary antibodies at 4°C. Cells were subsequently washed 3X with PBS and incubated with secondary antibodies diluted in blocking solution for 1.5 hour at RT, followed by 3X PBS washes. For fluorophore conjugated secondary antibodies, samples were individually treated with the specific conjugated antibodies for 2 hours at RT after secondary antibody incubation, followed by 3X washes with PBS. Finally, cells were counterstained using Hoechst 33343 (Invitrogen #H3570) in PBS for 20 min followed by 3X PBS washes. For wheat germ agglutinin (WGA) staining, cells were treated with WGA conjugated to Alexa Fluor 555 (Thermo Fisher #W32464) prior to permeabilization for 10 min at RT. Fresh PBS was added after the final wash to the cells and imaged on an LSM880 high resolution confocal microscope with a 63X objective using AiryScan imaging. The following primary antibodies were used: mouse anti-human GPHN (Synaptic Systems #147011); rabbit anti-human SOX10 (ThermoFisher #PA5-84795); rabbit anti-human KRT17 conjugated to Alexa Fluor 546 (Santa Cruz Biotechnology #sc-393002 AF546); rabbit anti-GABA (Sigma #A2052). The following secondary antibodies were used: AlexaFluor 488 anti-mouse IgG (Cell Signaling #4408S); AlexaFluor 647 anti-rabbit IgG (Cell Signaling #4414S).

#### Immunofluorescence staining of patient samples

Malignant melanoma in situ (MMIS) samples and Tumor microarrays (TMA) were obtained from the Memorial Sloan Kettering Cancer Center TMA Database and US Biomax (ME208). Paraffin sections of all tumors and normal skin were deparaffinized and antigen retrieval was performed using heat induced epitope retrieval method (HIER) with 1X Antigen Retrieval solution (Invitrogen #00-4955-58) in a pressure cooker at 95 for 10 min. Slides were washed 3X with PBS and blocked with 10% normal goat serum (ThermoFisher #50062Z) for 2 hours at RT and incubated with primary antibodies overnight at 4. Slides were washed with PBS (3X) followed by incubation with secondary antibodies for 2 hours at RT and 3X washes with PBS post-secondary antibody incubation. For fluorophore conjugated secondary antibodies, samples were individually treated with the specific conjugated antibodies for 2 hours at RT after secondary antibody incubation, followed by 3X washes with PBS. After the final PBS wash, slides were counterstained with Hoechst 33342 (Invitrogen #H3570) and mounted in ProLong Glass Antifade Mountant (Fisher #P36984). The following primary antibodies were used: rabbit anti-human GAD1 (Sigma #HPA058412) mouse anti-human GPHN (Synaptic Systems #147011); rabbit anti-human KRT17 (Sigma #HPA000452); rabbit anti-human S100A6 conjugated to Alexa Fluor 647 (Abcam #ab204028). The following secondary antibodies were used: AlexaFluor 488 anti-mouse IgG (Cell Signaling #4408S); Alexa Fluor 555 anti-rabbit IgG (Cell Signaling #4413S). For calculating GAD1 IF score, we probed the intensity (score 0-2) and the coverage (score 1-4) of each individual sample from melanoma patients and normal skin and assigned the IF score by multiplying the intensity score with the coverage score. All sections were imaged on an LSM880 high resolution confocal microscope with a 63X objective using AiryScan imaging.

### Proliferation assays

#### A375 phospho-Histone H3 (pH3) immunostaining

For proliferation studies, A375 melanoma cells (Azurite fluorophore positive) were co-cultured either with switched (GFP positive) or parental (dsRED positive) keratinocytes in low serum (2%) media, where melanoma cells and keratinocytes were plated in a 1:5 ratio for 48 hours. Following the 48 hours incubation, cells were fixed in 4% PFA and stained with a phospho-Histone H3 primary antibody (1:1000; Millipore #5-806) overnightat 4°C. AlexaFluor 647 anti-mouse IgG (Cell Signaling #4410) and more than 10 images per condition were acquired in a Zeiss AxioObserver fluorescence microscope. The number of mitotic cells was quantified by calculating double-positive cells (Azurite+, AlexaFluor647+) as a fraction of the total number of Azurite+ cells in each field.

#### CellTiter-Glo assay in melanoma cells treated with keratinocyte conditioned media

For conditioned media experiments, switched and parental keratinocytes were grown in 2% serum containing media for 24 hours and collected in 50 ml Falcon tubes followed by centrifugation at 500g for 5 minutes to remove any dead cells or debris. To measure cell proliferation, monocultures of A375 azurite positive cells were plated at a density of 5,000 cells per well in a 96-well plate in ioned media from switched or parental keratinocytes. After 48 hours of conditioned media treatment, CellTiter-Glo reagent (Promega #G7570) was added to the cells as per the manufacturer’s instructions and luminescence was read using a BioTek Synergy 96-well plate reader. All values were normalized to the A375 cells treated with parental keratinocytes conditioned media, done in quadruplicate for all cell lines. For the antagonist treatments, the same protocol as above was used except cells were treated with the following antagonists which were directly added to the conditioned media: AZD4547 (FGF receptor, Fisher Scientific #NC0660421, 100 ng/ml) or DMH1 (BMP receptor, Sigma #D8946, 0.5 µM) or EC330 (LIF receptor, Fisher #501871773, 30 nM).

#### Cyquant proliferation assay in monocultures and co-cultures

For proliferation experiments upon treatment with GABA/picrotoxin, 2000 or 5000 cells/well of melanoma cells were plated in 96-well plates. 24 hours post plating, GABA/picrotoxin was added to the media followed by a 48 hour incubation. Proliferation was measured using the Cyquant Cell Proliferation assay as per the manufacturer’s instructions and fluorescence was read using a BioTek Synergy 96-well plate reader. All values were normalized to the control conditions.

### Transmission Electron Microscopy

Melanoma/keratinocyte co-cultures were incubated for 48 hours and fixed using 4% paraformaldehyde, 2.5% glutaraldehyde, 0.002% picric acid in 0.1 mol/L sodium cacodylate buffer, pH 7.3 for transmission electron microscopy (12,000–15,000x). Imaging was performed on the JEOL JSM 1400, operated at 100 kV. Images were captured on a Veleta 2K × 2K CCD camera (EM-SIS).

### Electrophysiology studies

#### MEA assay

Extracellular recordings – Melanoma cell and keratinocyte co-cultures were seeded onto complementary metal oxide semiconductor multi-electrode array (CMOS-MEA) probes (3Brain AG, Wädenswil, Switzerland) coated with poly D-lysine (Sigma #P6407) overnight in the 37°C incubator. A 100-μl droplet of cell suspension was placed directly on the probe recording area. After 1 hour of incubation at 37°C, 1.5 ml of medium was added to the probe and replaced daily. Recordings were performed 48 hours days after plating. 2 minutes of spontaneous activity were sampled from the 4096 electrodes in each probe using the BioCAM X system. Recordings were analyzed using the BrainWave 4 software. Spike detection was performed using a Precise Timing Spike Detection (PTSD)^93^ algorithm by applying a threshold of 8 standard deviation to each channel trace. Average firing rates were calculated from a subset containing the 64 most active electrodes in each recording.

#### Calcium activity assay

Calcium activity was measured using the Cal-520 (Abcam ab171868) and Rhod-4 (Abcam ab112156) dyes following the manufacturer’s instructions. Briefly, melanoma and keratinocyte monocultures and co-cultures were plated in white bottom 96-well plates (Fisher #07-200-566) and incubated for 48 hours. Following incubation, cells were treated with Cal-520 or Rhod-4 dyes at 37 followed by RT incubation. For Cal-520 measurements, Picrotoxin (Sigma #P1675) was added to monocultures and co-cultures following RT incubation. Readings were measured using a BioTek Synergy 96-well plate reader at 488 and 540 nm wavelengths respectively.

### Data analysis

#### TCGA data analysis in melanoma patients

Melanoma patient sample analysis was performed using data from 448 samples in the TCGA Skin cutaneous melanoma (SKCM) PanCancer Atlas obtained from the Cancer Genome Atlas (TCGA)^94^. Samples were divided into two groups from gene expression data based on GAD1 high and GAD1 low mRNA expression. Kaplan Meier survival curves were generated using Graphpad Prism 9.

#### Analysis of publicly available GAD1 expression data

For GAD1 expression analysis, microarray expression data from Wistar melanoma cell lines was downloaded from Rockland-inc.com Melanoma Cell Lines and Mutations dataset (34 lines listed) (https://rockland-inc.com/melanoma-cell-lines.aspx)^33,95^. GAD1 expression value for each melanoma cell line (radial growth phase, vertical growth phase and metastatic) and normal cell lines (melanocytes, keratinocytes and fibroblasts) was plotted and GAD1 positivity assigned based on default analysis parameters used in the dataset.

### Supplementary materials

Table S1 (LOPAC Screen results in melanoma keratinocyte co-cultures)

Table S2 (RNA-seq and GSEA results in switched vs non-switched keratinocytes)

